# Spatial, temporal and numerical regulation of polar flagella assembly in *Pseudomonas putida*

**DOI:** 10.1101/2023.12.15.571843

**Authors:** Marta Pulido-Sánchez, Antonio Leal-Morales, Aroa López-Sánchez, Felipe Cava, Fernando Govantes

## Abstract

The Gram-negative bacterium *Pseudomonas putida* bears a tuft of flagella at a single cell pole. New flagella must be assembled *de novo* every cell cycle to secure motility of both daughter cells. Here we show that the coordinated action of FimV, FlhF and FleN sets the location, timing and number of flagella assembled. The polar landmark proteins FimV and FlhF are independently targeted to the nascent new pole during or shortly after cell division, but FimV stabilizes FlhF association with the cell poles. FlhF determines the polar position of the flagella by targeting early flagellar components to the cell pole and preventing their nucleation at non-polar sites. FlhF also promotes efficient flagellar assembly and indirectly stimulates Class III flagellar promoter activation by promoting secretion of the anti-FliA anti-σ factor FlgM. The MinD-like ATPase FleN partitions between the cell poles and the cytoplasm. Cytoplasmic FleN regulates flagellar number by preventing excessive accumulation of FlhF at the cell poles that may otherwise lead to hyperflagellation, likely by antagonizing FleQ-dependent transcriptional activation. FimV is essential to FleN polar location. FimV and FleN temporally regulate the onset of flagellar assembly by preventing premature polar targeting of FlhF and the ensuing premature targeting of additional flagellar components. Our results shed new light on the mechanisms that ensure the timely assembly of the appropriate number of flagella at the correct polar location in polarly flagellated bacteria.

**HIGHLIGHTS:**

- FimV, FlhF and FleN determine the position, number and timing of flagellar assembly
- FimV is essential to the normal intracellular distribution of FlhF and FleN
- FlhF restricts flagellar location to the cell poles and promotes efficient assembly
- Soluble, cytoplasmic FleN prevents polar FlhF accumulation and hyperflagellation
- Pole-bound FimV and FleN prevent premature FlhF recruitment and flagellar assembly

**GRAPHICAL ABSTRACT:** 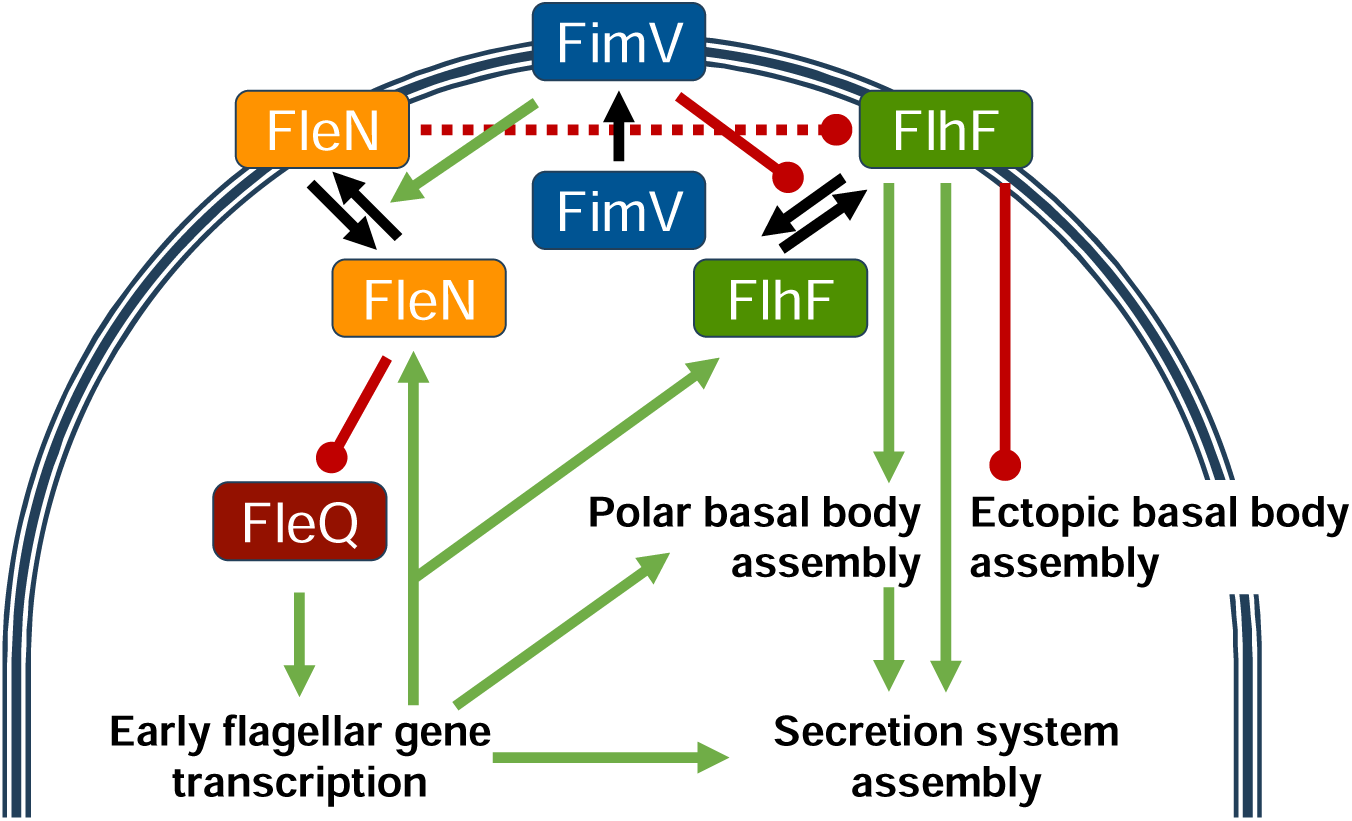

## INTRODUCTION

For many bacteria, the planktonic stage is hallmarked by the synthesis of one or more flagella, intricate molecular engines that enable motility in a liquid or semi-solid medium by engaging in propeller-like rotation (Altegoer *et al.,* 2014). Flagella are made of three functional units (**Fig. 1**): the basal body, a rotary machine embedded in the cell envelope; the filament, a long and thin helical structure that protrudes from the cells and acts as a propeller; and the hook, a flexible, curved element that connects basal body and filament. The basal body is in turn made out of five structural elements: the membrane-bound MS-ring lends support to the rest of components; the cytoplasmic C-ring modulates the direction of flagellar rotation; the flagellar type III secretion system (fT3SS) exports the extracellular flagellar components; the rod is a hollow cylinder that acts as a drive shaft for rotation and connects the MS-ring and the hook; and the cell wall-bound LP-ring surrounds the rod and plays the role of bushing. The flagellar stator, a membrane-bound complex that couples proton transport to the generation of torque is associated to the periphery of the basal body (Chevance & Hughes, 2008; Terashima *et al*., 2008). Flagellar function is modulated by the chemotaxis complexes, containing specific chemoreceptors and signal transduction proteins that interact with the C-ring to promote directional motion in response to concentration gradients (Matilla & Krell, 2017). The flagellar apparatus is sequentially assembled from the inside out (Chevance & Hughes, 2008), and flagellar genes are regulated by transcriptional cascades whose logic mirrors this assembly hierarchy (Smith & Hoover, 2009). Exquisite regulation of this process is paramount, as the synthesis and operation cost of the flagellar/chemotaxis system represents a substantial fraction of the energy resources of the cell (Macnab, 1996; Martínez-García *et al*., 2014).

**Figure 1.**
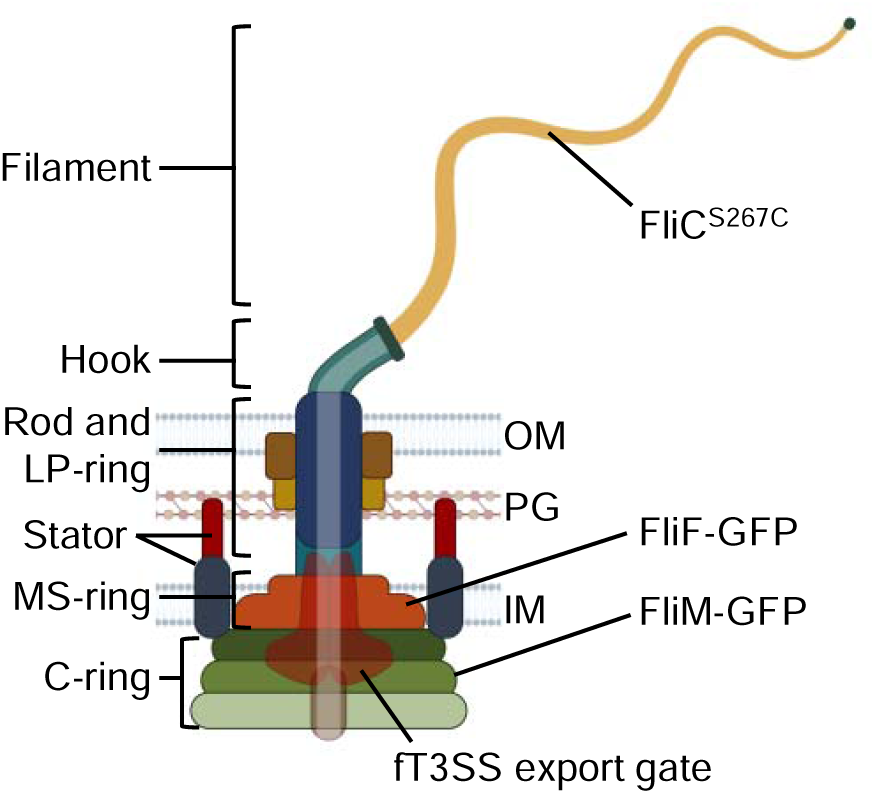
Structure of the Gram-negative flagellum. Cartoon depicting the archetypical structural units of the flagellum in Gram-negative bacteria and the location of the fluorescent flagellar protein derivatives used in this work: FliF-GFP (MS-ring), FliM-GFP (C-ring) and FliC^S267C^ (filament). OM: outer membrane; PG: peptidoglycan; IM: inner membrane; fT3SS: flagellar type III secretion system. Not drawn to scale. Created in part with Biorender.com.

Bacterial flagellation patterns are diverse and genetically determined in a species-specific fashion. Peritrichous flagella are located along the bacterial cell. Polar flagella may occur as single units in one (monotrichous) or both poles (amphitrichous), or as tufts containing a variable number of flagella (lophotrichous) in one or both poles. Less commonly, bacterial species naturally bearing their flagella in lateral or subpolar positions have also been described (Leifson, 1951; Schuhmacher *et al*., 2015). In peritrichously flagellated bacteria, such as *Escherichia coli*, flagella are assembled around the cell body and cell division results in a more or less even distribution of the existing flagella between both daughter cells to allow effective motility (Schuhmacher *et al*., 2015). In contrast, in bacteria displaying an unipolar flagellation pattern only one of the daughter cells inherits the preexisting flagellar machinery after cell division, and a new set of flagella must be assembled at the opposite pole to support motility of the other daughter cell (Amako & Umeda, 1982). To this end, cells must deploy specific regulatory mechanisms to ensure the correct location and number of flagella and precise timing to match the emergence of the new functional flagella with the completion of cell division. Much like chromosome replication and segregation, the synthesis and assembly of unipolar flagella can be envisioned as a discontinuous process that must be initiated in each cell cycle and terminated concomitantly to cell division.

*Pseudomonas putida* is a well-characterized Gram-negative soil bacterium endowed with great metabolic versatility and a model organism for biodegradation of organic toxicants and bioremediation (Martins dos Santos *et al*., 2004). During planktonic growth, *P. putida* displays unipolar lophotrichous flagellation. Most flagellar and core chemotaxis genes are located in a single chromosomal region, known as the flagellar cluster. Our recent work has unveiled the transcriptional organization of the flagellar cluster and the regulation of the flagellar gene set by a three-tiered regulatory cascade (Leal-Morales *et al*., 2022). FleQ is the master regulator of the flagellar/chemotaxis system in *P. putida*. FleQ activates σ^54^-dependent Class II flagellar promoters to drive the synthesis of all the structural components of the basal body and hook, and part of the core chemotaxis apparatus. FleQ also activates the synthesis of the alternative σ factor FliA. FliA is directly accountable for the activation of all the remaining (Class III) flagellar promoters (Leal-Morales *et al*., 2021), and several chemoreceptor-encoding genes (Rodríguez-Herva *et al*., 2010).

The *P. putida* flagellar cluster encodes at least two proteins potentially involved in the spatial, temporal and numerical regulation of flagellar synthesis. The MinD-like ATPase FleN (also known as FlhG or YlxH), and the SRP family GTPase FlhF are major players in determining the location and number of flagella in numerous bacterial species. FlhF is generally considered as a landmark protein that determines polar location of the flagella, while FleN is a negative regulator that limits the number of flagella (reviewed by Kazmierczak & Hendrixson, 2013; Altegoer *et al*., 2014; Schuhmacher *et al*., 2015). Our recent work has shown that deletion of *flhF* in *P. putida* results in the assembly of one or two flagella at random positions, while deletion of *fleN* leads to hyperflagellated cells with a unipolar lophotrichous pattern (Navarrete *et al*., 2019). FleN is also an antagonist of FleQ activation of the Class II flagellar promoters and *fleN* deletion provokes the overexpression of multiple flagellar genes (Navarrete *et al*., 2019; Leal-Morales *et al*., 2022).

*P. putida* genomes also encode orthologs of FimV, a *Pseudomonas aeruginosa* polar landmark protein. *P. aeruginosa* FimV promotes polar assembly of type IV pili (Carter *et al*., 2017), contributes to the activation of virulence genes *via* cAMP-mediated signaling (Inclan *et al*., 2016) and regulates the planktonic to biofilm transition (Bense *et al*., 2019; Schniederberend *et al*., 2019; Nicastro *et al*., 2020). FimV-related proteins, designated FimV, HubP or TspA, are present in multiple species of β- and γ-proteobacteria (Buensuceso *et al*., 2016). Members of the FimV/HubP/TspA family share a conserved architecture consisting of a N-terminal periplasmic domain, a single transmembrane region and a C-terminal cytoplasmic domain, but sequence conservation is limited to a periplasmic peptidoglycan-binding LysM motif (**IPR018392**) and a ∼50 residue distal C-terminal motif known as the FimV motif (**IPR020011**) (Wehbi *et al*., 2011; Buensuceso *et al*., 2017). HubP proteins from *Vibrio* spp. *and Shewanella putrefaciens* are polar landmarks involved in flagellar and chemotaxis complex biogenesis and replication origin segregation among others (Yamaichi *et al*., 2012; Rossmann *et al*., 2015 Takekawa *et al*., 2016; Arroyo-Pérez & Ringgaard, 2021; Altinoglu *et al*., 2022; Rick *et al*., 2022).

Here we demonstrate the involvement of FlhF, FleN and FimV in the temporal, spatial and numerical regulation of flagellar biogenesis in *P. putida*. Our results suggest that the polar recruitment of FimV and FlhF shortly after cell division and the dynamic association of FleN with the cell poles are required for the timely assembly of the correct number of flagella at the appropriate polar location. These findings shed light on the complex regulatory mechanisms required to ensure the faithful inheritance of the motile phenotype by both daughter cells in unipolarly flagellated bacteria.

## MATERIALS AND METHODS

### Bacterial strains and growth conditions

Bacterial strains used in this work are listed in **Supplemental Table S1**. Liquid cultures of *E. coli* and *P. putida* were routinely grown in Luria-Bertani (LB) medium (Sambrook & Russell, 2000) at 37 °C and 30 °C, respectively, with 180 rpm shaking. For solid media, American Bacteriological Agar (Condalab) was added to a final concentration of 15 g l^-1^. When needed, antibiotics and other compounds were added at the following concentrations: ampicillin (100 mg l^-1^), carbenicillin (500 mg l^-1^), chloramphenicol (15 mg l^-1^), rifampicin (20 mg l^-1^), gentamycin (10 mg l^-1^), kanamycin (25 mg l^-1^), 3-methylbenzoate (3MB) (3 mM) and sodium salicylate (2 mM). All reagents were purchased from Sigma-Aldrich.

### Plasmid and strain construction

Plasmids and oligonucleotides used in this work are summarized in **Supplemental Table S1.** All DNA manipulations were performed following standard protocols (Sambrook & Russell, 2000). Restriction and modification enzymes were used following to the manufacturer’s instructions (New England Biolabs). PCR amplifications were carried out using Q5^®^ High-Fidelity DNA Polymerase (New England Biolabs) for cloning and DreamTaq™ DNA Polymerase (Thermo Fisher) to verify plasmid constructions and chromosomal manipulations. One-step isothermal dsDNA Gibson assembly was performed as described (Gibson, 2011). *E. coli* DH5α was used as host strain for cloning procedures. Cloning steps involving PCR were verified by commercial Sanger sequencing (Stab Vida). Plasmid DNA was transferred to *E. coli* strains by transformation (Inoue *et al*., 1990), and to *P. putida* by triparental mating (Espinosa-Urgel *et al*., 2000) or electroporation (Choi *et al*., 2006). Site-specific integration of miniTn7-derivatives in *P. putida* strains was performed as previously described (Choi *et al*., 2005), and PCR-verified. Specific details of plasmid and strain construction are provided as **Supplemental Materials and Methods**.

### Swimming motility assays

Swimming assays were adapted from Parkinson (1976). Fresh colonies were toothpick-inoculated in Tryptone-agar plates (1 % Bacto^TM^ Tryptone [Difco], 0.5 % NaCl, 0.3 % Bacto^TM^ Agar), and incubated at 30 °C for 12 h. Digital images were taken and swimming halos diameter were measured using GIMP v2.10.12 (https://www.gimp.org) and normalized to the wild-type. At least, three biological replicates were assayed for each strain.

### *In vivo* gene expression assays

End-point fluorescence measurements were taken to quantify gene expression in strains harboring *gfp*-*lacZ* transcriptional fusions to different promoters. Overnight LB cultures supplemented with the corresponding antibiotics were diluted in 3 mL of the same medium to an A_600_ of 0.01 and incubated at 30 °C with 180 rpm shaking until reaching A_600_ of 0.4-0.6 (exponential phase). Samples of 1 mL were withdrawn and cells were washed three times in phosphate-buffered saline (PBS). Cells were resuspended in 500 µL of PBS and 150 µL were dispensed by triplicate into the wells of Costar^®^ 96-well polystyrene plates (Corning). A_600_ and GFP fluorescence (485 nm excitation, 535 nm emission, gain value set to 55) were measured in a Tecan Spark 10M microplate reader. Three biological replicates were assayed for each strain.

### RNA isolation and qRT-PCR

Total RNA from exponential phase cultures (A_600_ of 0.4) was extracted as described (García-González *et al*., 2005). Reverse transcription (RT) of 2 µg of total RNA into cDNA was carried out using the High-Capacity cDNA Reverse Transcription kit (Applied Biosystems) following the manufacturer’s instructions. Transcript levels of selected genes were quantified by real-time quantitative PCR (qPCR) using a CFX Connect Real-Time PCR Detection System (Bio-Rad), using appropriate oligonucleotides detailed in **Supplemental Table S1**. Reactions were prepared using the FastGene^®^ IC Green qPCR Universal kit (Nippon Genetics) following the manufacturer’s instructions. Specificity of the oligonucleotides was checked by the single-peak melting curve. Three biological replicates were assayed in triplicate.

### Protein secretion assay

Overnight cultures of strains expressing FlgM-FLAG were 100-fold diluted in 50 mL of LB medium supplemented with 2 mM salicylate and grown to A_600_ of 4. Cells were harvested by centrifugation (5000 *g*, 4 °C, 15 min) and culture supernatants were collected for further precipitation. To increase the detection threshold in the intracellular fraction, pelleted cells were resuspended in 5 mL of cold buffer B (20 mM Tris-HCl pH 8.0, 150 mM NaCl, 0.5 % Nonidet P-40) and disrupted by sonication. Soluble fractions were rotating-incubated overnight at 4 °C with 20 µL of buffer-equilibrated Anti-FLAG^®^ M2 magnetic beads (Sigma-Aldrich), followed by 10 washes with 1 mL of the same buffer. Culture supernatants were filtered through Millipore membranes with 0.45 µM pore-size (Merck) and TCA was added to a final concentration of 10 % (v/v). After overnight incubation at 4 °C, precipitates were collected by centrifugation (8000 *g*, 4 °C, 30 min) and washed 3 times with 1 mL cold acetone. Protein pellets were dried for 10 min at 100 °C. Precipitated supernatants and immunoprecipitated soluble cell fractions were further analyzed by Western-Blot.

### SDS-PAGE and Western-Blot analysis

Cell pellets and protein samples were mixed with 2x loading buffer (150 mM Tris-HCl pH 6.8, 20 % glycerol, 4% SDS, 0.1 % bromophenol blue, 10 % β-mercaptoethanol) and boiled at 100 °C for 10 min. Samples were centrifuged at 18000 *g* for 1 min and supernatants were separated on 12.5 % SDS-PAGE gels. Proteins were transferred to nitrocellulose membranes using a Trans-Blot Turbo Transfer System (Bio Rad). Protein transference was evaluated by Ponceau S Staining Solution (Thermo Fischer). Membrane was blocked for 2 h at room temperature in blocking solution (3 % fat-free milk in TBS buffer with 1 % Tween 20). Blocked membrane was incubated overnight at 4 °C with primary antibody anti-FLAG (F3165, Sigma-Aldrich, 1:2000 dilution) or anti-GFP (SAB4301138, Sigma-Aldrich, 1:5000 dilution), washed 3 times in TBS with 1% Tween 20 (TBS-T), probed for 2 h at 4 °C with secondary antibody anti-mouse (31430, Thermo Fischer, 1:10000 dilution) or anti-rabbit (31460, Thermo Fischer, 1:10000 dilution) HRP-conjugated, and washed with TBS-T as before. Membranes were added Pierce™ ECL Western Blotting Substrate (Thermo Fischer) and chemiluminescent signal was revealed using a ChemiDoc MP instrument (Bio-Rad) with a exposure time of 800 s. Image Lab software (Bio-Rad) was employed for image analysis.

### Brightfield microscopy videos

Continuous videos of near-surface motility were recorded using a Leica DMI4000B inverted microscope equipped with a Leica HCX PL FLUOTAR 20x/0.40 CORR PH1 objective lens and a Leica DFC360 FX CCD high speed camera (pixel size: 0.2015 μm). Overnight LB cultures were 250-fold diluted in the same medium and grown to early-exponential phase (A_600_ of 0.2-0.3). Sample volumes of 150 μl of serial dilutions 10^3^ to 10^5^ were transferred to the wells of a Costar^®^ 96-well polystyrene plate (Corning) and brightfield videos of cells swimming on the plane of the well surface were recorded for 1 min.

### Time-lapse fluorescence microscopy

Overnight cultures were 100-fold diluted in LB medium and grown to early-exponential phase (A_600_ of 0.2-0.3). Culture drops (2 µL) were placed on LB-agarose pads (1 % [w/v] agarose in 10 % [v/v] LB, to minimize background fluorescence) in glass slides and cells were imaged every 5 min using an Axio Imager.Z2 (Zeiss) microscope equipped with a Plan-Apochromat 63x/1.40 Oil Ph 3 M27 contrast objective lens and an ORCA-Flash 4.0 LT digital CMOS camera (Hamamatsu, pixel size: 0.1031 μm), controlled by the Zeiss ZEN 2 (Blue edition) software enabling software autofocus for focus maintenance over time. Phase contrast and GFPmut3 fluorescence images were respectively acquired using a TL Vis-LED Lamp (4.91 Volt, exposure time of 50 ms) and the LED module 470 nm from a Colibri.2 LED light source using a Zeiss 38 HE Green Fluorescent Prot. filter (excitation filter BP: 450/490 nm, emission filter BP: 500/550 nm, exposure time of 500 ms).

### Confocal microscopy

Overnight cultures were 100-fold diluted in LB medium supplemented with 2 mM salicylate when necessary and grown to exponential phase (A_600_ of 0.4-0.6). Cells from 1 mL samples were harvested by centrifugation at 5000 *g* for 5 min and washed three times in PBS to remove LB. Flagellar filaments of strains expressing FliC^S267C^ were stained by gently resuspending cells in 100 µL of PBS containing 5 µg/mL Alexa Fluor^TM^ 488 C_5_ maleimide (Molecular Probes), followed by incubation for 5 min at room temperature in the dark (Hintsche *et al*., 2017). Cells were washed three times for excess dye removal. For membrane staining, cells were resuspended in 100 µL of PBS containing 5 µg/mL FM™ 4-64 (Molecular Probes). After incubation for 5 min at room temperature in the dark, 2 µL drops were immediately placed on 1 % agarose pads made with deionized water to be imaged using an Axio Observer7 confocal microscope (Zeiss) equipped with a CSU-W1 spinning disk module (Yokogawa). Images were taken using a Zeiss α Plan-Apochromat 100x/1.46 Oil DIC M27 objective lens and a Prime 95B CMOS camera (Teledyne Photometrics, pixel size: 0.1099 μm), controlled by the SlideBook 6 software. GFPmut3 and Alexa Fluor^TM^ 488-labeled FliC^S267C^ fluorescence was excited with the 488 nm laser line and collected with emission filter 525/50 nm, while FM™ 4-64 was excited at 561 nm and collected with a 617/73 filter (exposure time of 1000 ms). Exposure times for GFPmut3-tagged FimV, FlhF, FliF and FliM were 300 ms, 1000 ms for FleN and 100 ms for Alexa Fluor^TM^ 488-labeled FliC^S267C^. For each image, seven Z-sections were collected with a step size of 0.33 μm.

### Microscopy image analysis

Microscopy images processing and analysis were performed using Fiji software v1.53t (Schneider *et al*., 2015). Brightfield microscopy continuous videos were converted to 4 fps time-lapses and trajectories of 20 cells of each strain were tracked for 15 s using Manual Tracking v2.1.1 plugin bundled with Fiji. Background subtraction of the fluorescence signal from GFPmut3, Alexa Fluor^TM^ 488 C_5_ maleimide and FM™ 4-64 was set to a rolling ball radius of 20 pixels. Confocal images are shown as maxima projections of 7 Z-sections of the 525/50 channel, merged with the 617/73 channel showing the focal plane with the cell contour. Cell segmentation, maxima and foci detection, and cell measurements from confocal images were performed using MicrobeJ v5.13I plugin (Ducret *et al*., 2016). Unless otherwise stated, default parameters were applied. Cell segmentation was performed on the lowest Z-plane of the brightfield channel using the rod-shaped descriptor (area: 1-9 µm^2^, length: 2-5.5 µm, width: 0.5-3 µm). The old cell pole was defined as the one showing the highest maxima intensity from the 525/50 channel. Images showing the same fluorescence-labeled protein were batched-processed using the same maxima/foci tolerance and Z-score parameters, with a minimal intensity cutoff value of 200 and foci area of 0.02-max µm^2^. A Gaussian-fitting algorithm was applied to discriminate between two close foci (radius: 1 pixel, method: oval, iterations: 1000, minimal r^2^ for optimization: 0.9). A tolerance distance of 0.3 µm outside the segmented cell boundary was set for maxima/foci association to parent cells. Analyzed images were manually inspected to confirm that the majority of cells were properly assigned. Shape descriptors, medial axis profiles, polarity, maxima/foci intensities and parent localization associations were recorded for each strain. Demographic representations of the fluorescence intensity of the 525/50 channel along the cell medial axis were performed for 500 randomized cells of each strain, sorted by cell length. Fluorescent signal at midcell was manually inspected. MicrobeJ analysis-derived data were collected from three independent sets of 500 randomized cells within datasets containing at least 2000 segmented cells for each strain. Alexa Fluor^TM^ 488-labeled FliC^S267C^ filaments were manually inspected in 500 randomized cells from sets of 8 images containing 100-200 segmented cells for each strain.

### Statistical analysis

Statistical treatment of the data was performed using GraphPad Prism v8.3.0 software. Results are reported as the average and standard deviation of at least three biological replicates or image datasets, or median and 95% confidence interval for the median for fluorescence foci quantification. Significance of the differences among strains was evaluated by means of the two-tailed Student’s t-test for unpaired samples not assuming equal variance.

### Bioinformatics analyses

Sequence alignments were performed using the MUSCLE package at the EMBL-EBI web site (https://www.ebi.ac.uk/Tools/msa/muscle/) (Edgar, 2004). Similarity between FimV/HubP proteins was scored using BLASTP at the National Center for Biotechnology Information web site (https://blast.ncbi.nlm.nih.gov/Blast.cgi) (Altschul *et al*., 1990). Operon prediction was performed using the Operon-mapper web server (https://biocomputo.ibt.unam.mx/operon_mapper/) (Taboada *et al.,* 2018).

## RESULTS

### FimV is required for full flagellar function

The *P. putida* KT2440 ORF PP_1993 encodes an ortholog of *P. aeruginosa* polar landmark protein FimV. *P. putida* FimV displays 55% identity and 68% similarity to its *P. aeruginosa* counterpart. To assess the possible involvement of FimV in flagellar motility we constructed MRB97, a derivative of *P. putida* KT2442 (a spontaneous rifampicin-resistant mutant of the reference strain KT2440) bearing a complete deletion of PP_1993. The Δ*fimV* mutant displayed 39%-reduced motility halo diameter relative to the wild-type in a soft agar-based swimming assay, while a non-motile Δ*fleQ* mutant, used here as negative control, did not spread from the inoculation point (**Fig 2A**). We also used time-lapse phase contrast microscopy to analyze the near-surface swimming trajectories of the wild-type and Δ*fimV* strains (**Fig. 2B**). Compared to the wild-type, most Δ*fimV* cells displayed shorter, erratic trajectories indicative of slower swimming speed. These results indicate that FimV is not essential to flagellar motility, but is required for normal flagellar function.

**Figure 2.**
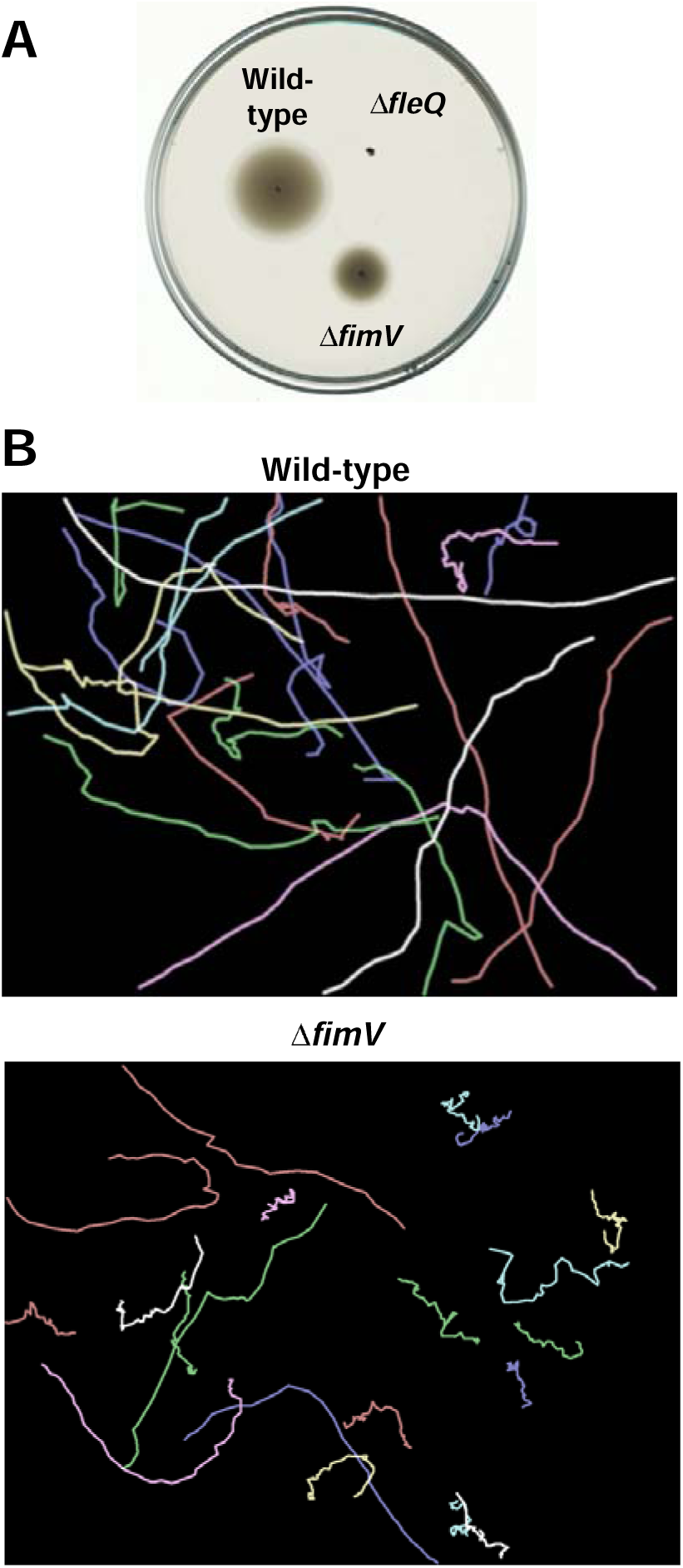
Swimming motility of the Δ*fimV* mutant. **A.** Soft agar-based swimming assay of the Δ*fimV* mutant. The wild-type strain and the Δ*fleQ* mutant assayed in the same plate were used as positive and negative controls. The picture shows a representative swim plate out of at least three separate replicates. **B.** Near-surface swimming trajectories of wild-type and Δ*fimV* cells. Each coloured line represents the trajectory of a cell tracked for 15 s from brightfield microscopy videos (n=20 cells).

### FimV and FlhF accumulate at the new cell pole during or shortly after cell division

Our previous work showed that FlhF is essential to the polar location of flagella in *P. putida*, and a Δ*flhF* mutant also displayed reduced swimming motility and short swimming trajectories (Navarrete *et al*., 2019). We assessed the intracellular location of FimV and FlhF by expressing fluorescent C-terminal GFP fusions from miniTn*7* derivatives inserted at the chromosomal *att*Tn*7* site of the wild-type *P. putida* strain KT2442. The FlhF-GFP fusion was expressed from its physiological P*flhF* promoter (Navarrete *et al*., 2019) and translational start signals. On the basis of bioinformatic operon predictions (**Taboada *et al*., 2018**), we expressed the FimV-GFP fusion from its upstream gene *asd* promoter region (P*asd*) and *fimV* natural translation initiation signals. The fluorescent fusion proteins fully complemented the swimming defects of the corresponding deletion mutants (**Fig. S1 A and B**), and qRT-PCR revealed that *fimV* transcript levels produced from P*asd* were similar to those obtained from the native *fimV* locus of the wild-type strain (**Fig. S2A**).

FimV and FlhF formed fluorescent foci at one or both poles in the vast majority (93%) of wild-type cells, with 33 and 34% displaying a unipolar and 60 and 58% a bipolar distribution, respectively (**Fig. 3A and B**). Bipolar distribution *vs.* cell length plots derived from demographic analysis (**Figs. S3 and Fig. 3C**) showed that ∼50% of the shorter newborn cells bore bipolar foci of FimV-GFP and FlhF-GFP. The frequency increased with cell length to reach a maximum of 85-90% in cells longer than ∼3.5 µm. Fluorescent FimV-GFP and FlhF-GFP foci were also found at the midcell region in 8-9% of the cells. Demographic analysis showed that midcell foci were restricted to cells longer than ∼3.8 µm, being present in almost 50% of the longer, predivisional cells (**Fig. 3D and E**). Midcell foci were often associated to an incipient constriction indicative of cell division, and the fluorescent signal occasionally appeared as a transversal band spanning the width of the cell (**Fig. S4**). We explored the dynamics of FimV-GFP and FlhF-GFP association with the cell poles further by time-lapse fluorescence microscopy imaging in the wild-type strain. This experiment was performed with FimV produced from the salicylate-inducible P*sal* promoter. In the absence of inducer, this construct complemented the motility defect of the Δ*fimV* mutant and displayed a very similar localization behavior (**Fig. S5 A-F**), although it produced 6- to 7-fold more *fimV-gfp* transcript and fusion protein (**Fig. S2A and B**). Salicylate induction was deleterious to *P. putida* growth and was not used in further experiments. The time-lapse images showed most cells displaying FimV-GFP foci at both poles (**Fig. 4A and Supplemental movie S1**). New foci emerged at the midcell region of dividing cells displaying visible constrictions, or at the new cell poles shortly after cell division, and remained associated the cell poles for the duration of the experiment (300 minutes). A similar dynamics was obtained with the FlhF-GFP fusion (**Fig. 4B and Supplemental movie S2**). Taken together, our results suggest that FimV and FlhF are recruited to the emerging new poles concomitant to or shortly after cell division and remain permanently bound to the cell poles. This repeated cycle results in the predominantly bipolar distribution observed for both proteins.

**Figure 3.**
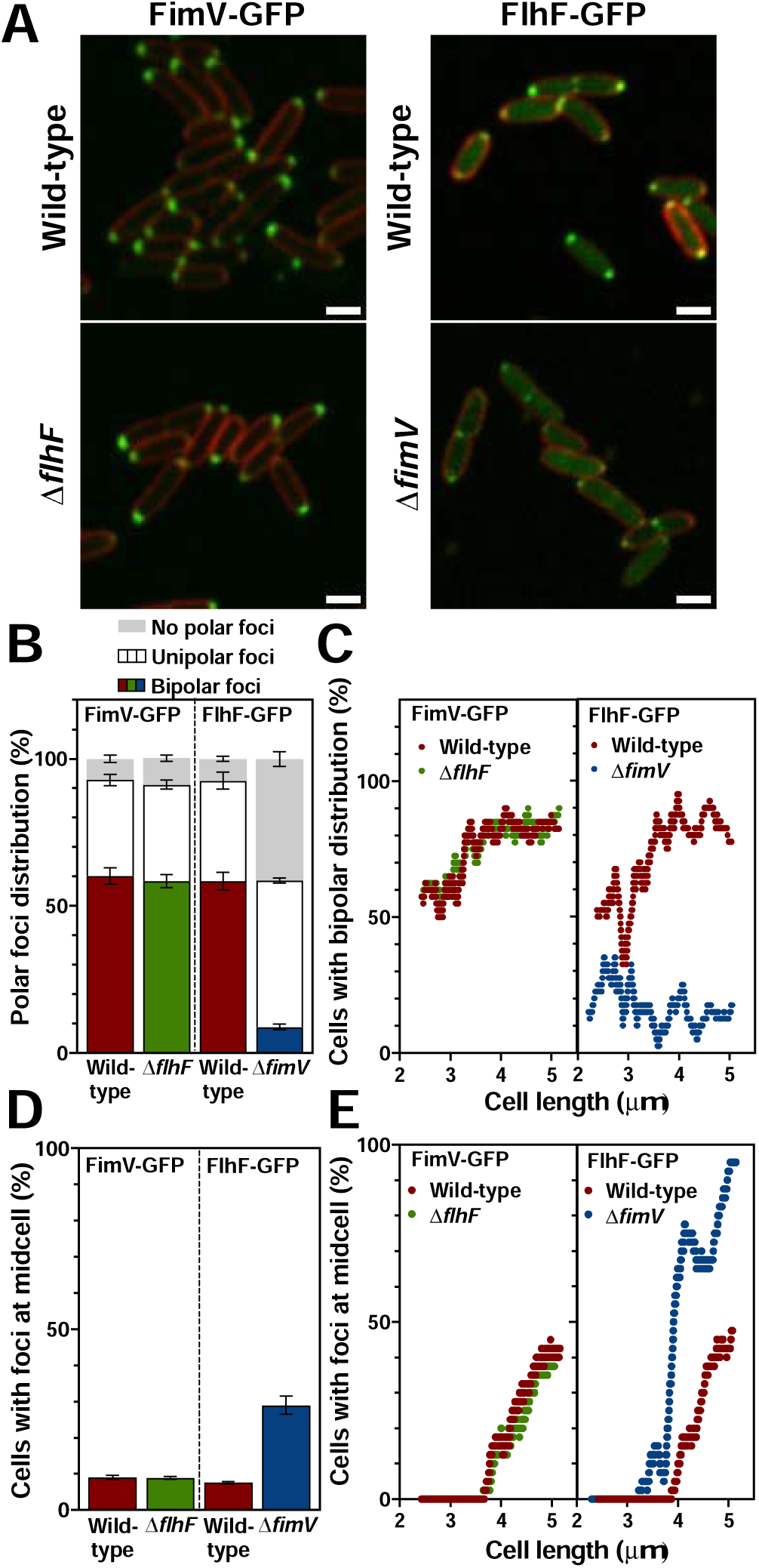
Intracellular location of FlhF and FimV. **A.** Confocal microscopy images of wild-type, Δ*flhF* and Δ*fimV* cells expressing P*asd-fimV-gfp* or P*flhF-flhF-gfp* (green). FM^TM^ 4-64 was used as membrane stain (red). Images are shown as the maxima projections of seven Z-sections of the green channel, merged with the red channel showing the cell contour at the focal plane. Scale bar: 2 µm. **B.** Frequency of wild-type, Δ*flhF* and Δ*fimV* cells bearing bipolar, unipolar or no polar FimV-GFP or FlhF-GFP foci. **C.** Frequency of wild-type, Δ*flhF* and Δ*fimV* cells displaying bipolar distribution of FimV-GFP or FlhF-GFP foci *vs*. cell length (n=500 cells). **D and E.** Frequency of wild-type, Δ*flhF* and Δ*fimV* cells displaying midcell FimV-GFP or FlhF-GFP foci (**D**) and the same parameter plotted against cell length (**E**) (n=500 cells). Columns and error bars represent averages and standard deviations of at least three separate replicates. Stars denote *p-*values of the two-tailed Student’s T test not assuming equal variance (*=*p*<0.05; **= *p*<0.01; ***= *p*<0.001; ****= *p*<0.0001).

**Figure 4.**
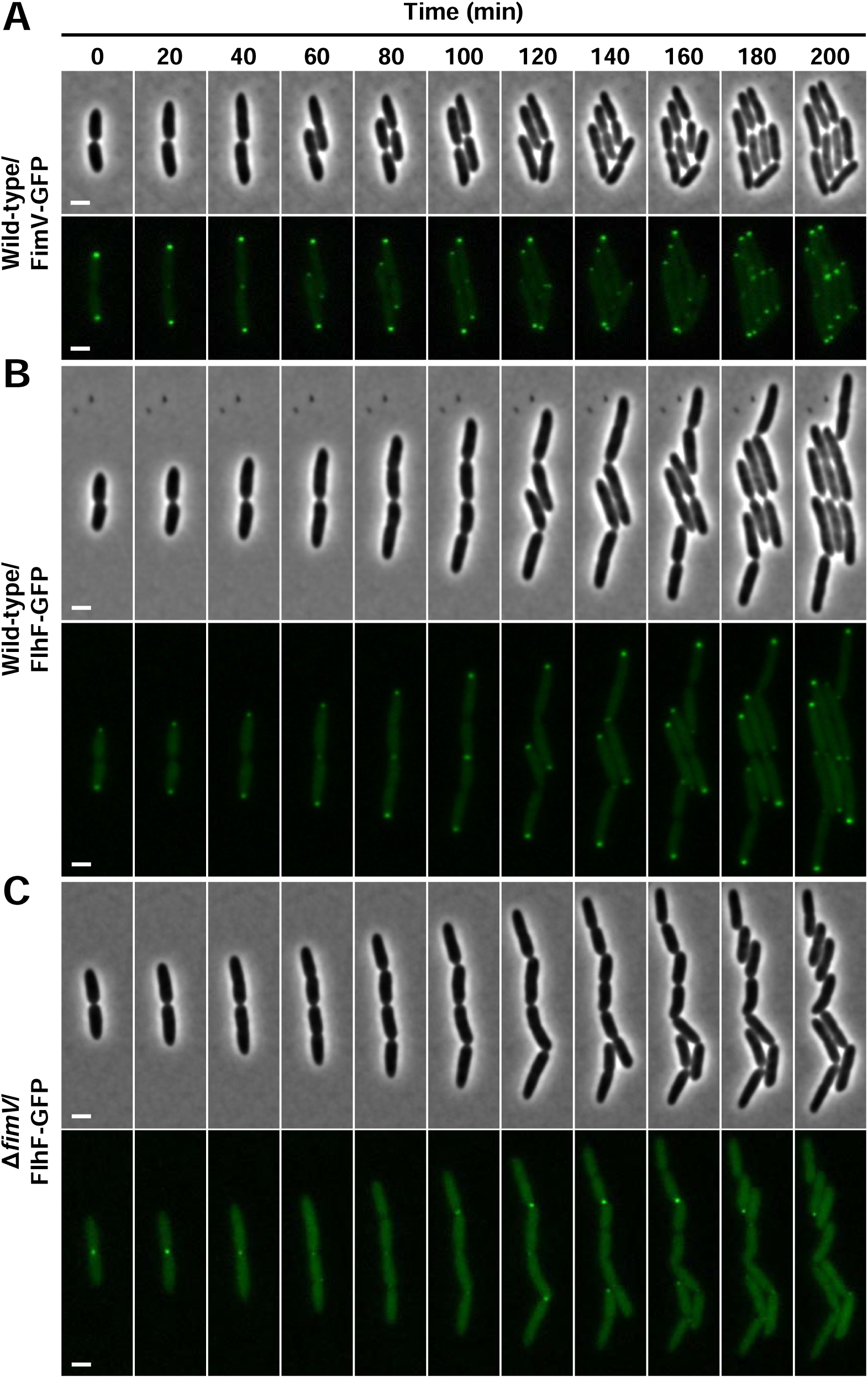
Intracellular distribution dynamics of FimV and FlhF. Time-lapse fluorescence microscopy images of wild-type cells expressing P*sal-fimV-gfp* (**A**) or wild-type (**B**) and Δ*fimV* cells (**C**) expressing P*flhF-flhF-gfp*. Phase contrast (odd rows) and fluorescence (even rows) images are shown. Scale bar: 2 µm.

### FimV stabilizes FlhF association with the cell poles

The intracellular distribution of FlhF-GFP was deeply disturbed in the Δ*fimV* background. Visual inspection and quantitative analysis revealed a clear decrease in unipolar and bipolar FlhF-GFP distribution in the Δ*fimV* mutant resulting in a large proportion of vacant poles (**Fig 3A and 3B**). In addition, the frequency of bipolar distribution no longer increased with cell length. Instead, a maximum frequency of ∼30% was found in short (∼2.5 µm) cells that decreased with length to 10-15% in cells longer than 4.5 µm. However, midcell foci were more prevalent, being present in over 90% of the longer cells, suggesting that recruitment to the nascent new poles occurs earlier in the absence of FimV (**Fig. 3D and E**). To test the contribution of FimV to FlhF recruitment further, we monitored the dynamics of FlhF-GFP accumulation at the cell poles of the Δ*fimV* mutant by time-lapse fluorescence microscopy (**Fig 4C and Supplemental movie S3**). Deletion of *fimV* did not prevent FlhF-GFP recruitment to the new pole, but newly formed foci were stochastically disassembled (average lifespan of ∼75 minutes) in the absence of FimV (**Fig. S6**). These results suggest that polar targeting of FlhF does not require FimV, but FimV promotes a stable association of FlhF with the cell poles. Similarly, we assessed the impact of FlhF on FimV-GFP location. Our results showed no significant alteration in any of the parameters measured (**Fig. 3A-E**), suggesting that FlhF has no influence of the intracellular distribution of FimV.

### FlhF restricts flagellar assembly to the cell poles

We next questioned the impact of FlhF and FimV on the cellular localization of the structural components of the flagella. To this end, we engineered FliF and FliM protein fusions to GFP, along with the flagellin FliC derivative FliC^S267C^, bearing a serine-to-cysteine substitution to enable binding of fluorescent maleimide-derived fluorochromes. FliF, FliM and FliC are structural components of the MS- and C-rings and the filament, respectively (**Fig. 1**). All three proteins were produced in mini-Tn*7* derivatives from their natural promoters (P*fliE*, P*fliL* and P*fliC*) and translational start signals and were shown to complement the null swimming phenotypes of the corresponding deletion mutants (**Fig. S7**). Analysis of FliF-GFP, FliM-GFP and FliC^S267C^ intracellular distribution in the wild-type strain revealed the presence of polar FliF-GFP and FliM-GFP foci and FliC^S267C^ filaments in >90% of the population (**Figs. 5A and B**). Bipolar foci or filaments were found in 48, 28 and 2% of the cells (**Fig. 5B**), and demographic analysis revealed that bipolar foci accumulated with cell growth reflecting the expected order of assembly, with FliF preceding FliM and FliC filaments emerging significantly only in the longer cells (**Figs. S8A and B and 5C**). Non-polar FliF-GFP and FliM-GFP foci were only found in 8 and 6% of the cells, while non-polar filaments were almost non-existent (**Fig. 5D**). These results suggest that *P. putida* flagellar structures are assembled efficiently and specifically at the new cell pole during the course of a cell cycle in preparation for filament emergence during the following cell division.

**Figure 5.**
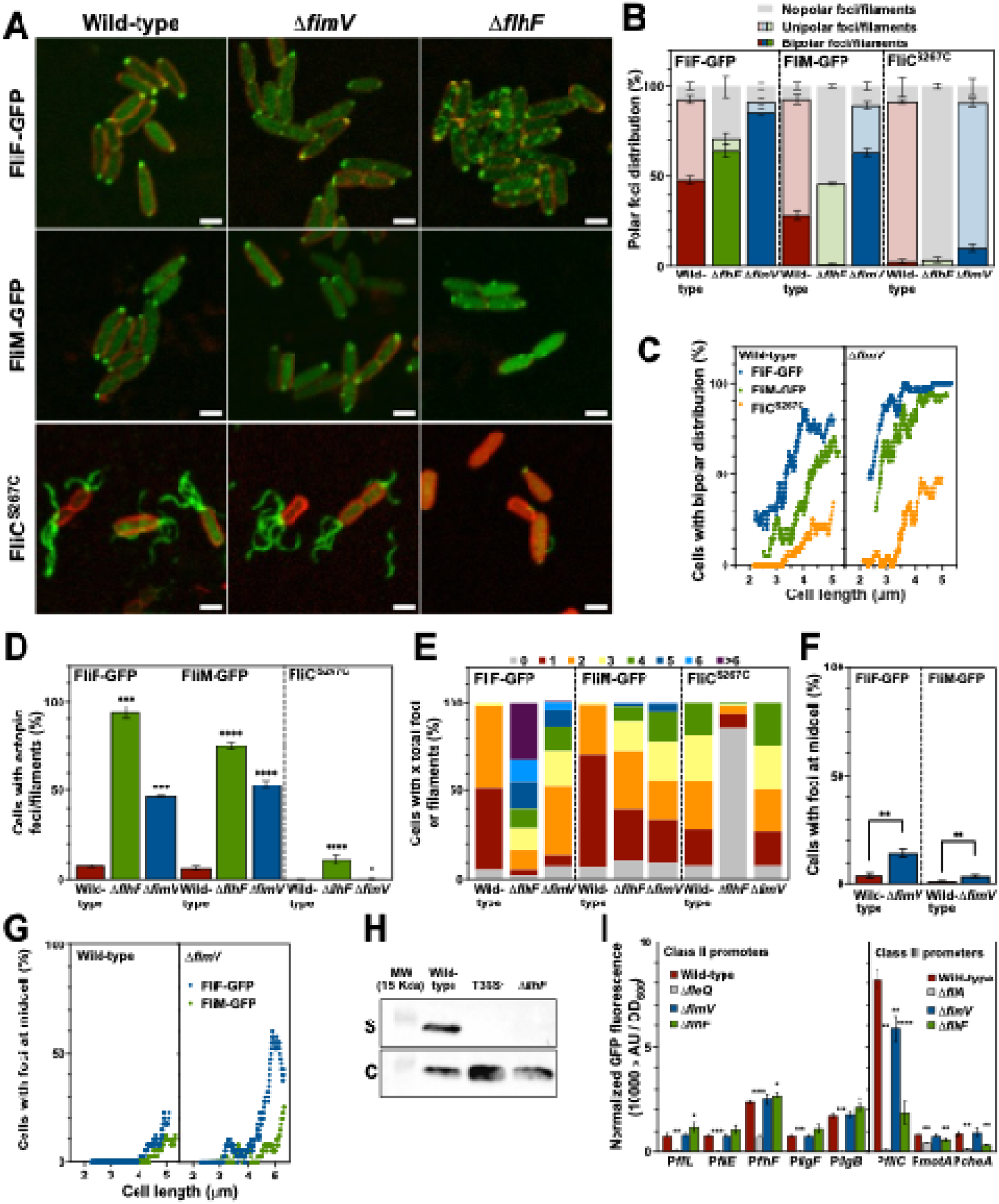
Analysis of the recruitment of flagellar structural proteins in the Δ*fimV* and Δ*flhF* mutants. **A.** Confocal microscopy images of wild-type, Δ*fimV* and Δ*flhF* cells expressing P*fliE-fliF-gfp*, P*fliL-fliM-gfp* or P*fliC-fliC*^S267C^ (green). Alexa Fluor^TM^ 488 C_5_ maleimide was used for FliC^S267C^ filament staining. FM^TM^ 4-64 was used as membrane stain (red). Images are shown as the maxima projections of seven Z-sections of the green channel, merged with the red channel showing the cell contour at the focal plane. Scale bar: 2 µm. **B.** Frequency of wild-type, Δ*flhF* and Δ*fimV* cells bearing bipolar, unipolar or no polar FliF-GFP or FliM-GFP foci, or FliC^S267C^ filaments. **C.** Frequency of wild-type, Δ*flhF* and Δ*fimV* cells displaying bipolar distribution of FliF-GFP or FliM-GFP foci, or FliC^S267C^ filaments *vs.* cell length (n=500 cells). **D.** Frequency of wild-type, Δ*flhF* and Δ*fimV* cells bearing non-polar FliF-GFP or FliM-GFP foci, or FliC^S267C^ filaments. **E.** Percentage of wild-type, Δ*flhF* and Δ*fimV* cells bearing 0, 1, 2, 3, 4, 5, 6 or more than 6 total FliF-GFP or FliM-GFP foci, or FliC^S267C^ filaments. **F and G.** Frequency of wild-type and Δ*fimV* cells displaying midcell FliF-GFP or FliM-GFP foci (**F**) and the same parameter plotted against cell length (**G**) (n=500 cells). **H.** Western blot analysis of FLAG-tagged FlgM in the culture supernatants (S) and cell (C) fractions of the wild-type, fT3SS^-^ and Δ*flhF* strains. MW: molecular weight marker (15 kDa band shown). **I.** Normalized GFP fluorescence from exponential phase cultures of wild-type, Δ*fleQ,* Δ*fliA,* Δ*fimV* and Δ*flhF* strains bearing transcriptional P*fliL-gfp*, P*fliE-gfp*, P*flhF-gfp*, P*flgF-gfp*, P*flgB-gfp*, P*fliC-gfp*, P*motA-gfp* or P*cheA-gfp* fusions. Columns and error bars represent averages and standard deviations of at least three separate replicates. Stars denote *p-*values of the two-tailed Student’s T test not assuming equal variance (*=*p*<0.05; **= *p*<0.01; ***= *p*<0.001; ****= *p*<0.0001).

In the absence of FlhF, FliF-GFP and FliM-GFP foci were no longer restricted to the cell poles (**Fig. 5A**), as evidenced by a decrease in the frequency of polar foci (**Fig. 5B**), and a sharp increase in the frequency of non-polar foci (**Fig. 5D**). Indeed, the vast majority of Δ*flhF* cells displayed multiple FliF-GFP foci (median = 5) and at least one FliM-GFP focus (median = 2) at apparently random locations (**Fig. 5A and E**). Complementarily, we assessed the assembly hierarchy of the basal body to show that FliF-GFP can form polar foci in the absence of FliM or in the presence of a transposon insertion that precludes transcription of the fT3SS components *fliP*, *fliQ*, *fliR* and *flhB* (fT3SS^-^ mutant) (**Fig. S9A and B**). In contrast, polar location of FliM-GFP is strictly dependent on the presence of FliF and partially on the fT3SS (**Fig. S9B**). Accordingly, the presence of FliM-GFP foci is a sign that incipient flagellar assembly incorporating at least FliF and FliM occurs in the vast majority of Δ*flhF* cells. In contrast, FliC^S267C^ filaments were only observed in ∼15% of the Δ*flhF* cells. Most of these cells displayed only one or two unusually short filaments, often less than one wavelength long, in polar or non-polar positions (**Figs. 5A and E and S10**), consistent with previously published flagellar stain images (Navarrete *et al*., 2019). The infrequent occurrence of FliC^S267C^ filaments relative to FliM-GFP or FliF-GFP foci suggests that in the absence of FlhF the efficiency of flagellar assembly declines sharply as it progresses from the incorporation of basal body proteins FliF to FliM to the flagellin FliC. Taken together, our results indicate that FlhF restricts flagellar location to the cell pole and fosters efficient assembly of the flagellar apparatus.

Despite the impact of FimV on FlhF localization, the phenotypes of the Δ*flhF* mutant were not replicated by the Δ*fimV* mutant. Instead, polar FliF-GFP and FliM-GFP foci and FliC^S267C^ filaments were present in the vast majority of cells (**Figs. 5A and B**) and the numbers of filaments per cell were similar to those in the wild-type (**Fig. 5E**), indicating that an apparently normal polar flagellar complement is assembled in the absence of FimV. The frequency of cells with bipolar distribution was higher for all three proteins and demographic analysis revealed that bipolar foci accumulated at shorter cell lengths (**Fig. 5B and C**), indicating that, similarly to FlhF, the structural flagellar proteins are recruited earlier in the absence of FimV. Frequent non-polar FliF-GFP and FliM-GFP foci and occasional non-polar FliC^S267C^ filaments were also detected in this background (**Figs. 5D and E**), suggesting that unstable association of FlhF with the cell poles in the absence of FimV is sufficient to promote efficient polar flagella assembly, but cannot prevent incipient flagella assembly at non-polar locations. As shown above for FlhF (**Fig. 3E**), midcell FliF-GFP and FliM-GFP also appeared with low frequency in the wild-type strain, and their frequency was increased in the Δ*fimV* mutant up to ∼25 and ∼50% in longer cells, respectively (**Fig. 5F and G**). These results suggest that premature recruitment of FlhF in the absence of FimV promotes early assembly of the flagellar components

### FlhF is required for fT3SS-dependent secretion and full Class III flagellar gene expression

We have shown above that most Δ*flhF* cells produce flagellar assembly intermediates containing FliF and FliM at both polar or non-polar locations that rarely progress into complete flagella. Since most flagellar components beyond the cell membrane are secreted by way of the fT3SS, we hypothesized that flagellar secretion may be impaired in the absence of FlhF. To test this possibility, we used FlgM, the FliA-specific anti-σ factor, as a reporter of fT3SS-dependent secretion. To this end, we produced a FlgM-FLAG fusion from the induced P*sal* promoter in a mini-Tn*7* derivative and detected it in cell fractions and culture supernatants by Western blot using an anti-FLAG antibody in the wild-type strain and the Δ*flhF* mutant (**Fig 5H**). The fT3SS^-^ mutant was used as a negative control. A band compatible with the molecular weight of FlgM-FLAG (∼12 kDa) was detected in the cell fractions of the three strains and in the wild-type strain supernatant, but was conspicuously absent in those of the fT3SS^-^ and Δ*flhF* mutants, indicating that FlgM is secreted by way of the fT3SS in *P. putida*, and FlhF is required for fT3SS function. We tested the functional implications of faulty FlgM secretion in the Δ*flhF* mutant by assessing the expression of a set of flagellar promoters fused to *gfp* (**Fig. 5I**). The five Class II promoters tested were positively regulated by FleQ as described (Leal-Morales *et al*., 2022), but only marginally upregulated in the Δ*flhF* mutant. The Class III P*fliC*, P*motA* and P*cheA* promoters were FliA-dependent as described (Leal-Morales *et al*., 2022), and also displayed 5-, 3- and 2-fold decreased expression in the Δ*flhF* mutant, suggesting that limited FlgM secretion in the absence of FlhF precludes FliA activation. In contrast, deletion of *fimV* did not have a major impact on any of the Class II or Class III promoters, consistent with the normal assembly of a full complement of polar flagella observed in this strain.

### FimV is required for polar location of FleN

We have previously shown that FleN is a negative regulator of the number of flagella in *P. putida*. Accordingly, a Δ*fleN* mutant produces an unusually high number of polar flagella, is defective in swimming motility and displays abnormal swimming trajectories (Navarrete *et al*., 2019). To assess the intracellular distribution of FleN, C-terminal and N-terminal fusions of *fleN* to *gfp* expressed from P*flhF* were constructed, but failed to complement the swimming defect of the Δ*fleN* mutant. An N-terminal FleN-GFP fusion expressed from the salicylate-inducible P*sal* promoter only complemented slightly the swimming defect of the Δ*fleN* when expressed in the presence and in the absence of inducer (**Supplemental Fig. S11**), suggesting that the fusion protein retains partial function. Accordingly, we used this partially active N-terminal GFP-FleN fusion in this study. Because P*sal* induction did not improve complementation, salicylate addition was omitted.

The GFP-FleN protein displayed a similar intracellular distribution to that observed with FimV-GFP and FlhF-GFP in the wild-type strain (**Fig. 6A and B**). Polar foci occurred in 86% of the cells, with 22% and 64% showing unipolar and bipolar distribution respectively. However, other features were divergent from this conserved behavior. Firstly, a high level of background fluorescence was present in the cell cytoplasm (**Fig. 6A**). Secondly, the bipolar distribution frequencies derived from demographic analysis did not correlate with cell length; instead, ∼60% of the cells bore bipolar GFP-FleN foci across the complete cell length scale (**Fig. 6C**). Finally, ectopic foci at diverse non-polar locations were present in 42% of the cells (**Fig. 6D**). These results indicate that, unlike FimV and FlhF, FleN partitions between the cell poles, the bulk cytoplasmic compartment and, to a lesser extent, other cellular locations independently of the stage in the cell cycle

**Figure 6.**
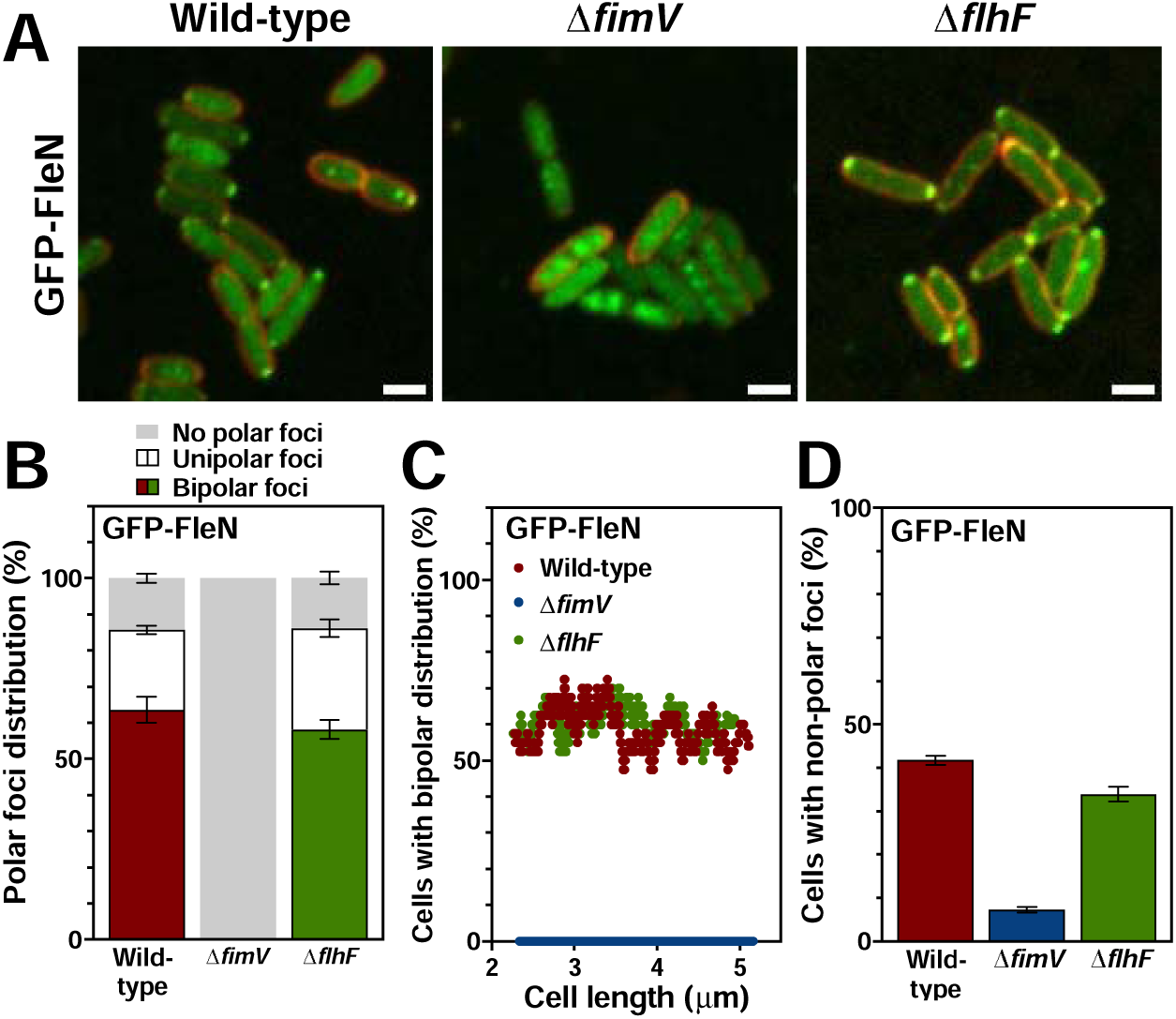
Intracellular location of FleN. **A.** Confocal microscopy images of wild-type, Δ*fimV* and Δ*flhF* cells expressing P*sal-gfp-fleN* (green). FM^TM^ 4-64 was used as membrane stain (red). Images are shown as the maxima projections of seven Z-sections of the green channel, merged with the red channel showing the cell contour at the focal plane. Scale bar: 2 µm. **B.** Frequency of wild-type, Δ*fimV* and Δ*flhF* cells bearing bipolar, unipolar or no polar GFP-FleN foci. **C.** Frequency of wild-type, Δ*fimV* and Δ*flhF* cells bearing bipolar, unipolar or no polar GFP-FleN foci *vs.* cell length (n=500 cells). **D.** Frequency of wild-type, Δ*fimV* and Δ*flhF* cells bearing ectopic GFP-FleN foci. Columns and error bars represent averages and standard deviations of at least three separate replicates. Stars denote *p-*values of the two-tailed Student’s T test not assuming equal variance (*=*p*<0.05; **= *p*<0.01; ***= *p*<0.001; ****= *p*<0.0001).

The intracellular distribution of GFP-FleN was radically altered by the deletion of *fimV* **Fig. 6A-D**). GFP-FleN was no longer associated to the cell poles (**Fig. 6A-C**) and the frequency of non-polar foci was also strongly reduced (**Fig. 6D**). Instead, the fluorescent signal was irregularly distributed in the cytoplasm (**Fig. 6A**). These results indicate that FimV function is essential to polar location of FleN. In contrast, GFP-FleN distribution was largely unaltered by deletion of *flhF* (**Fig. 6A-D**).

### FimV and FleN regulate the timing of early flagellar assembly

Next we examined the effect of FleN on the location of FimV and FlhF. While FimV-GFP intracellular distribution was not altered in the absence of FleN (**Fig. S12**), deletion of *fleN* caused an increase in the frequency of bipolar FlhF-GFP at both poles (**Fig. 7A and B**), and the demographic analysis revealed an increase in bipolar distribution in shorter cells, indicative of earlier FlhF recruitment to the new poles (**Fig. S2 and 7C**). Similarly, *fleN* deletion also increased the frequency of midcell FlhF-GFP foci, which is indicative of recruitment at shorter cell lengths, and a greater frequency was also observed in longer cells (**Figs. 7D and E and S2**). Because FlhF is necessary for polar location of the flagellar components (**Fig. 5**), we questioned whether premature FlhF recruitment would impact the polar targeting dynamics of the structural flagellar proteins. The Δ*fleN* mutant displayed increased bipolar distribution frequencies of FliF-GFP and FliM-GFP (**Fig. 7A and B**), which correlated with new pole targeting at shorter cell lengths (**Figs. 7C**). The frequency of midcell FliF-GFP and FliM-GFP foci was also increased in the Δ*fleN* mutant (**Fig. 7D**), and midcell FliF-GFP emerged at shorter cell lengths (∼3.5 µm *vs.* ∼4 µm) and appeared with increased frequency in longer cells of the mutant strains (maximum ∼50% *vs.* ∼20%) (compare **Fig. 7E** with **Fig. 5G**). A similar trend was observed with FliM-GFP foci, but the differences were small and the results inconclusive. These phenotypes are reminiscent of those observed above with the Δ*fimV* mutant (**Fig. 5B-F**). Taken together, our results suggest that FimV and FleN prevent premature association of FlhF and at least some components of the flagellar structure to the emerging new cell poles.

**Figure 7.**
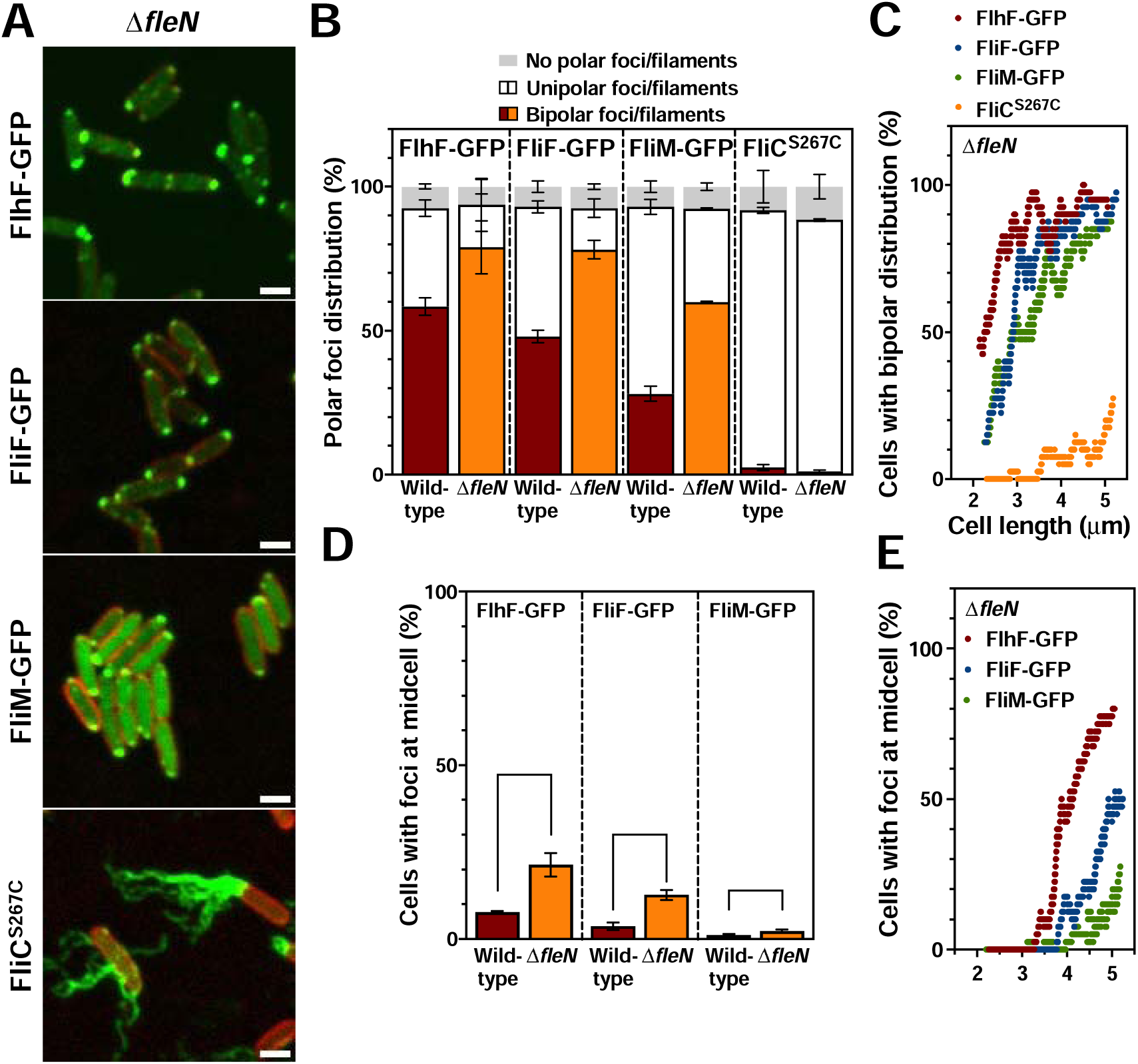
Cellular location of flagellar proteins in the Δ*fleN* mutant. **A.** Confocal microscopy images of wild-type and Δ*fleN* cells expressing P*flhF-flhF-gfp*, P*fliE-fliF-gfp*, P*fliL-fliM-gfp* or P*fliC-fliC^S267C^* (green). Alexa Fluor^TM^ 488 C_5_ maleimide was used for FliC^S267C^ filament staining. FM^TM^ 4-64 was used as membrane stain (red). Images are shown as the maxima projections of seven Z-sections of the green channel, merged with the red channel showing the cell contour at the focal plane. Scale bar: 2 µm. **B.** Frequency of wild-type and Δ*fleN* cells bearing bipolar, unipolar or no polar FlhF-GFP, FliF-GFP or FliM-GFP foci, or FliC^S267C^ filaments. **C.** Frequency of wild-type and Δ*fleN* cells displaying bipolar distribution of FlhF-GFP, FliF-GFP or FliM-GFP foci, or FliC^S267C^ filaments *vs.* cell length (n=500 cells). **D and E.** Frequency of wild-type and Δ*fleN* cells displaying midcell FlhF-GFP, FliF-GFP or FliM-GFP foci (**D**) and the same parameter plotted against cell length (**E**) (n=500 cells). Columns and error bars represent averages and standard deviations of at least three separate replicates. Stars denote *p-*values of the two-tailed Student’s T test not assuming equal variance (*=*p*<0.05; **= *p*<0.01; ***= *p*<0.001; ****= *p*<0.0001).

### FleN prevents polar accumulation of FlhF and the structural flagellar proteins

We have previously shown that deletion of FleN causes hyperflagellation in *P. putida* (Navarrete *et al*., 2019). Quantitative analysis of the fluorescence microscopy images revealed an increase in FlhF-GFP foci intensity in the absence of FleN, suggesting that FleN restricts the amount of FlhF associated to the cell poles (**Fig. 8A**). Fluorescence intensity of the polar FliF-GFP and FliM-GFP foci was also higher in the absence of FleN (**Fig. 8B**) and, although, we could not reliably measure fluorescence of the FliC^S267C^ filaments, filament counts revealed that *fleN* deletion caused a clear increase in the number of filaments: while ∼75% of the wild-type cells displayed one to three filaments and only ∼18% displayed more than three units, ∼75% of the Δ*fleN* cells exhibited more than three filaments, and at least nine filaments were detected in some cells (**Fig. 8C**). These results suggest that FleN-limited accumulation of FlhF results in limited recruitment of the structural flagellar proteins to effectively prevent the production of an excessive number of flagella that leads to faulty flagellar function (Navarrete *et al*., 2019). Despite the crucial role of FimV in FleN association to the cell pole, filament numbers in the Δ*fimV* mutant were also similar to those in the wild-type, as shown above (**Fig. 5E**), and a comparable increase in fluorescent foci intensity was not obtained with the Δ*fimV* mutant for any of the structural flagellar proteins (**Fig. 8B**). Very low foci intensity values were obtained with FlhF-GFP (**Fig. 8A**), consistent with premature dissociation of the FlhF-GFP foci in this background, as shown above (**Fig. 4C**). These results indicate that FleN association with the cell poles is not required for inhibition of excessive polar accumulation of FlhF and the structural flagellar proteins and therefore the soluble, the non-pole-bound form of FleN is likely responsible for this phenomenon.

**Figure 8.**
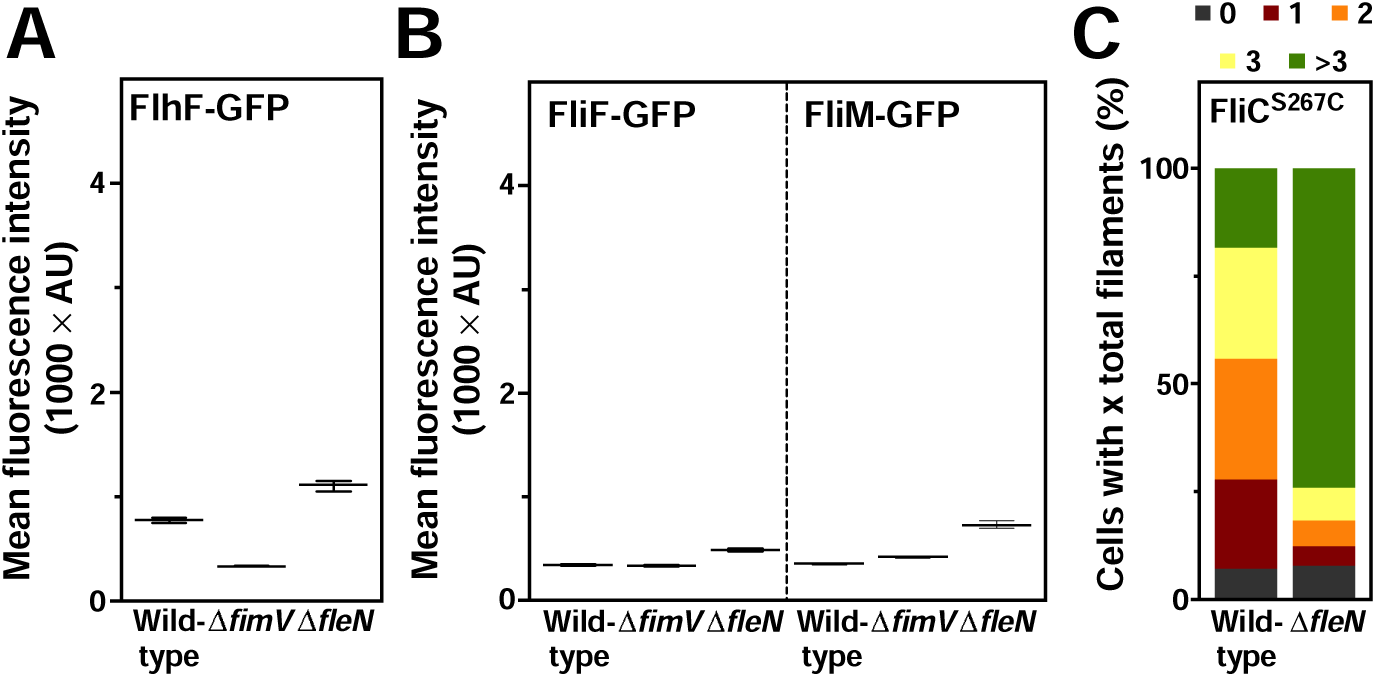
Accumulation of flagellar proteins in the Δ*fimV* and Δ*fleN* mutants. **A and B.** Mean fluorescence intensity of polar FlhF-GFP (**A**), FliF-GFP or FliM-GFP (**B**) foci in wild-type, Δ*fimV* or Δ*fleN* cells (n=500 cells). **C.** Percentage of wild-type and Δ*fleN* cells bearing 0, 1, 2, 3 or more than 3 total FliC^S267C^ filaments. Points correspond to individual fluorescence intensity values, bars denote the median and the 95% confidence interval for the median.

## DISCUSSION

Spatiotemporal and numerical control of flagellar assembly is a critical aspect of the regulation of motility in polarly flagellated bacteria (Kazmierczak & Hendrixson, 2013; Schuhmacher *et al*., 2015). The Gram-negative bacterium *P. putida* displays lophotrichous flagellation, bearing a tuft of flagella at a single pole. Because of the asymmetry of the unipolar flagellation pattern, motile *P. putida* cells can only transmit their flagellar machinery to one of their offspring upon cell division. Hence, to secure motility of both daughter cells, *P. putida* must (i) identify the new pole as a site for flagella assembly, (ii) assemble *de novo* a tuft with the correct number of flagella at this site, and (iii) coordinate the timing of such assembly with the cell cycle to ensure emergence of the new flagella and motility of the daughter cell after division. This work identifies and characterizes the involvement of a novel element, the polar landmark protein FimV, and expands on the function of two previously described elements, FlhF and FleN (Navarrete *et al*., 2019), in the spatiotemporal and numerical control of flagellar assembly in *P. putida*.

We have previously shown that deletion of *flhF* or *fleN* leads to defects in flagellar motility characterized by slower motility in soft agar-based assays and aberrant swimming trajectories (Navarrete *et al*., 2019). A Δ*fimV* mutant also displays motility defects in soft agar-based and microscopy analyses, consistent with a slower swimming speed (**Fig. 2**). Inactivation of the highly similar *P. aeruginosa* FimV did not decrease swimming (Buensuceso *et al*., 2017; Bense *et al*., 2019; Schniederberend *et al*., 2019; Nicastro *et al*., 2020). In contrast, inactivation of the more distantly related HubP proteins of *S. putrefaciens* and *Vibrio* spp. resulted in reduced swimming (Yamaichi *et al*., 2012; Rossmann *et al*., 2015; Takekawa *et al*., 2016; Park *et al*., 2019; Arroyo-Pérez & Ringgaard, 2021). Our results highlight the fact that members of the FimV/HubP/TspA family adopt flexible roles that may reflect the specific physiological needs of each host organism.

FimV is located at both cell poles in most *P. putida* cells (**Fig. 3A and B**), is recruited to the midcell region during cell division and remains associated to the cell poles (**Fig. 4A and Movie S1**). FimV-GFP accumulates at midcell in up to ∼50% of the longer cells, often associated with the visible constriction typically caused by Z-ring contraction (**Figs. 3E and S4**). However, a similar fraction of the longer cells does not accumulate FimV-GFP at midcell, and the frequency of bipolar foci increases from ∼50% to >80% during early growth after cell division (**Fig. 3C**), suggesting that recruitment to the new cell poles may also occur at this stage. *Vibrio cholerae* and *S. putrefaciens* HubP target the nascent poles during cell division and nucleate further at the new poles during cell growth (Yamaichi *et al*., 2012; Rossmann *et al*., 2015; Galli *et al*., 2017). Midcell recruitment of FimV or HubP to has been linked to cell division. On one hand, FimV binds peptidoglycan (Wehbi *et al*., 2011), and the peptidoglycan-binding LysM motif is essential to polar location in *S. putrefaciens* HubP and *P. aeruginosa* FimV (Rossmann *et al*., 2015; Buensuceso *et al*., 2017). Since septal peptidoglycan is targeted by proteins bearing LysM domains among others (Buist *et al*., 2008), FimV/HubP interaction with septal peptidoglycan may play a role in FimV recruitment or stabilization at the new cell pole (Wehbi *et al*., 2011). On the other hand, *V. cholerae*, HubP recruitment does not happen in the absence of FtsZ or FtsI, suggesting the involvement of the cell division machinery (Galli *et al*., 2017). In *P. putida* FimV-GFP appears as a transversal band spanning the cell width in cells bearing a minimal constriction (**Fig. S4**), reminiscent of the location of the septal ring. We propose that FimV recruitment may precede septal peptidoglycan synthesis, although interaction with septal/polar peptidoglycan may contribute to stabilize the interaction.

Similarly to FimV, FlhF displays a bipolar distribution in most wild-type cells (**Fig. 3A and B**), resulting from its recruitment to the nascent poles during or shortly after cell division and its stable association the cell poles (**Fig. 4B and Supplemental Movie S2**). FlhF is located at the cell membrane, associated to the flagellated cell pole or both cell poles in many polarly flagellated bacteria (Murray and Kazmierczak, 2006; Kusumoto *et al*., 2008; Green *et al*., 2009; Ainsaar *et al*., 2019; Arroyo-Pérez and Ringgaard, 2021). Initial FlhF recruitment is independent of FimV, but FimV is responsible for stable association with the cell poles (**Fig. 3A-E and 4B and Supplemental Movie S3**). FlhF recruitment is independent of HubP in *V.* spp. and *S. putrefaciens* (Yamaichi *et al*., 2012; Arroyo-Pérez and Ringgaard, 2021), but the frequency of polar location decreases notably in the absence of FimV in the latter (Rossmann *et al*., 2015). Polar location of FlhF is independent of any other flagellar proteins in *Vibrio* spp. and *Shewanella oneidensis* (Kusumoto *et al*., 2008; Green *et al*., 2009; Gao *et al*., 2015). Polar recruitment of FlhF and other proteins requires the action of the polar organelle coordinator (Poc) complex, comprised of homologs to the outer membrane transporter-energizing TonB-ExbB-ExbD complex, in *P. aeruginosa* and *P. putida* (Cowles *et al*., 2013; Ainsaar *et al*., 2019). The Poc complex is diffusely located at the cell periphery (Cowles *et al*., 2013), and the mechanism for polar protein recruitment is unclear. A recent preprint (Arroyo-Pérez *et al*., 2024) documents the involvement of FimV and FipA (formerly known as FlrD) in the recruitment of FlhF to the cell pole in *V. parahaemolyticus, P. putida, and S. putrefaciens*. In *P. putida*, deletion of *fimV* and *fipA* limit FlhF association to the cell poles in an additive fashion. FipA is recruited to the nascent new cell pole during or shortly after cell division, but dissociates during further cell growth. These results and our own can be reconciled in a model in which FimV, FipA and FlhF are targeted to the new cell pole early in the cell cycle, with FipA being essential to FlhF recruitment and FimV being responsible for permanent association of FlhF with the cell poles during cell growth.

FliF-GFP and FliM-GFP foci and FliC^S267C^ filaments were almost exclusively found at the wild-type cell poles (**Fig. 5A, B and D**). This strict polar location is obliterated in the Δ*flhF* mutant, where the three proteins could be found indistinctly at polar, non-polar locations or both (**Fig. 5A, B, D and E**). Delocalization of flagellar assembly in *flhF*-null mutants has been reported in *P. putida*, *P. aeruginosa* and *V. cholerae* (Pandza *et al*., 2000; Murray and Kazmierczak, 2006; Green *et al*., 2009; Navarrete *et al*., 2019). Because of the hierarchical nature of flagellar assembly, we propose that FlhF acts primarily by recruiting the early flagellar protein FliF to the cell pole. Pole-bound FliF may in turn guide specific polar assembly of the flagellar structure and prevent initiation of flagellar assembly elsewhere. This is supported by several observations: (i) FliF is the monomeric unit of the transmembrane MS-ring, which lends support to the complete supramolecular structure of the flagellum; (ii) FliF nucleates at both polar and non-polar locations in the absence of FlhF (**Figs. 5 A, B, D and E**); and (iii) polar recruitment of FliF does not require prior assembly of the fT3SS or the C-ring (**Fig. S9**). FlhF recruits FliF to initiate the flagellar assembly in *Vibrio* spp. (Green *et al*., 2009; Fukushima *et al*., 2023). Recent work in *S. putrefaciens* showed that a preformed complex comprising FlhF and the C-ring protein FliG is required to capture membrane-diffusing FliF proteins (Dornes *et al*., 2024). While our observations are fully compatible with both models, at this point we cannot discriminate if FlhF must be bound to FliG to recruit membrane-bound FliF.

Over 90% of the wild-type cells bore fluorescent FliF-GFP and FliM-GFP foci and FliC^S267C^ filaments in one pole (**Fig. 5B**). The high incidence of flagellated cells and the similar frequency of polar foci for all three proteins tested, suggest that flagellar assembly is highly efficient. Cooperative assembly of the basal body, rod and stator complexes and highly processive protein export *via* the fT3SS and filament assembly are documented mechanisms to ensure completion of the flagellar apparatus (Kubori *et al*., 1992; Minamino *et al*., 2014; Li and Sourjik, 2011; Minamino *et al*., 2021). Fluorescently labeled flagellar proteins are recruited to the new (non-flagellated) pole after FlhF and in the expected assembly order: FliF>FliM>FliC (**Figs. 3C and 5C**), consistent with the role of FlhF as initiator of flagellar assembly. We also note that less than 3% of the wild-type cells bear flagellar filaments at both poles. This outcome implies precise coupling of flagellar filament assembly with cell division to prevent both premature filament emergence before cell division resulting in bipolar flagellation and delayed filament emergence after cell division resulting in a significant number of aflagellate newborn cells.

Most Δ*flhF* mutant cells bore multiple FliF-GFP and a few FliM-GFP foci. Because FliM is a cytoplasmic protein displaying a diffuse cytoplasmic distribution in the absence of FliF (**Fig. S9**), we infer that, in the absence of FlhF, FliF can nucleate at random locations and recruit FliM, likely along with FliG, which is known to interact with both (Marykwas and Berg, 1996; Levenson *et al*., 2013; Dornes *et al*., 2024), to form incipient flagellar structures. Several lines of evidence suggest that the Δ*flhF* mutant does not support efficient flagellar assembly beyond these early complexes. Firstly, fT3SS-dependent secretion of FlgM was not detected in the absence of FlhF, suggesting that fT3SS activity is limited or absent. Secondly, FlhF is specifically required for full expression of the flagellar FliA-dependent Class III promoters, suggesting that FlgM secretion is an assembly checkpoint for late flagellar gene transcription, as previously described (Chevance & Hughes, 2008, Smith & Hoover, 2009). FlhF is required for an additional assembly checkpoint connecting early basal body assembly with the synthesis of late basal body and hook components activated by a dedicated two-component system (known as FleSR, FlrBC or FlgSR) in other polarly flagellated bacteria (Correa *et al*., 2005; Murray & Kazmierczak, 2006; Kim *et al*., 2012; Ren *et al*., 2018; Burnham *et al*., 2020). However, *P. putida* FleS and FleR are not required for flagellar component synthesis or activity (Leal-Morales *et al*., 2022), and late basal body and hook gene transcription is not downregulated in the absence of FlhF (**Fig. 5I**), indicating that this checkpoint does not operate in *P. putida*. Finally, only 15% of the Δ*flhF* cells displayed flagellar filaments, indicating that only a small fraction of the incipient FliF- and FliM-containing flagellar complexes lead to the assembly of complete flagella. Diminished frequencies of flagellated cells or complete loss of flagella have been reported in *flhF*-null mutants of multiple bacterial species (Pandza *et al*., 2000; Correa *et al*., 2005; Murray & Kazmierczak 2006; Salvetti *et al*., 2007; Kusumoto *et al*., 2008; Balaban *et al*., 2009; Kim *et al*., 2012), but the extent of assembly of early flagellar substructures has not been assessed. Flagellar filaments in the Δ*flhF* cells, when present, were few (generally, one or two), and notably shorter than those in the wild-type (**Figs. 5A and S10**), a likely consequence of poor FliC transcription in this background. We propose that FlhF not only restricts flagellar assembly to the cell poles, but also supervises the assembly process to ensure efficient progress and completion of a functional flagellar complement. In the absence of FlhF, frequent failure of the assembly process leads to a low rate of fT3SS secretion, compromising further incorporation of the secreted flagellar components, preventing release of FliA from FlgM-mediated inactivation, and ultimately restricting FliA-dependent gene expression.

The Δ*fimV* mutant displayed polar tufts with similar flagellar numbers and morphology to the wild-type (**Fig. 5A and E**), indicating that FimV-mediated stabilization of FlhF association with the cell pole is not required for the assembly of polar flagella. This is consistent with the fact that most often FlhF dissociation in the absence of FimV does not happen during the first cell cycle after recruitment (∼30 minutes in our experimental conditions) (**Fig. S6**). However, the Δ*fimV* mutant displayed frequent non-polar FliF- and FliM-containing assemblies that rarely led to filament completion (**Fig. 5C-E**), suggesting that permanent association with the cell poles appears crucial to prevent abortive flagellar assembly at ectopic positions.

FleN nucleates at the cell poles in most *P. putida* cells, but is also found diffusely distributed at the cell cytoplasm and forming additional foci at non-polar locations (**Fig. 6A, B and D**). While its *P. aeruginosa* and *S. oneidensis* orthologs were only found in the cell cytoplasm (Gao *et al*., 2015; Kawalek *et al*., 2023), those from *Vibrio* spp., *S. putrefaciens* and *Campylobacter jejuni*. display a tendency to partition between the cell poles and the cytoplasm (Kusumoto *et al*., 2008; Balaban and Hendrixson, 2012; Yamaichi *et al*., 2012; Blagotinsek *et al*., 2020). Polar targeting of FleN was strictly dependent on FimV (**Fig. 6**, **A-C**), as shown for HubP in *Vibrio* spp. (Yamaichi *et al*., 2012; Takekawa *et al*., 2016; Kojima *et al*., 2020; Arroyo-Pérez and Ringgaard, 2021). Unlike FimV and FlhF, FleN association with the cell pole was unaffected by cell division and growth (**Fig. 6C**). FlhG formed polar foci during cell division that were disassembled shortly after in *V. parahaemolyticus* (Arroyo-Pérez *et al*., 2021). FlhG polar localization relies on interactions with C-ring components FliM, FliN or FliY in *S. putrefaciens* and *C. jejuni* (Schuhmacher *et al*., 2015; Henderson *et al*., 2020; Blagotinsek *et al*., 2020). However, this is unlikely to be relevant to *P. putida*, where FleN location does not display the cell cycle-dependent dynamics characteristic of the structural flagellar proteins (**Figs. 5C and 6C**). We have not explored the involvement of ATP binding and hydrolysis on FleN location as previously described (Schuhmacher *et al*., 2015; Ono *et al*., 2015; Kojima *et al*., 2020).

FleN is a negative regulator that limits the number of flagella in *P. putida* (Navarrete *et al*., 2019). Increased number of flagella in the Δ*fleN* mutant is mirrored by increased polar accumulation of FlhF, FliF, FliM and FliC (**Figs. 7A and B and 8A-C**). FleN orthologs restrict the number of flagella to that characteristic of each species in the peritrichously flagellated *Bacillus subtilis* (Guttenplan et al., 2013; Schuhmacher, Rossmann, et al., 2015), the monotrichously flagellated *P. aeruginosa, S. oneidiensis* and *Vibrio* spp. (Correa *et al*., 2005; Dasgupta *et al*., 2000; Gao *et al*., 2015; Ono *et al*., 2015) and the amphitrichously flagellated *C. jejuni* (Gulbronson *et al*., 2016). Two mechanisms have been proposed to mediate the negative effect of FleN on the number of flagella. FleN is an auxiliary protein to the flagellar master regulator FleQ in *P. putida* and *P. aeruginosa*. FleN antagonizes FleQ activation of Class II flagellar genes and modulates FleQ regulation of other non-flagellar genes (Hickman and Harwood, 2008; Baraquet *et al*.,2012; Baraquet and Harwood, 2013; Navarrete *et al*., 2019; Leal-Morales *et al*., 2022). FleN orthologs negatively regulate flagellar gene expression in *V. cholerae* and *S. putrefaciens* (Correa *et al*., 2005; Blagotinsek *et al*., 2020). FlhF overproduction is sufficient to cause hyperflagellation in *P. putida* and other polarly flagellated bacteria (Pandza *et al*., 2000; Kusumoto *et al*., 2006; Green *et*lJ*al*., 2009; Schniederberend *et*lJ*al*., 2013), suggesting that structural flagellar components are synthesized in excess, and the number of flagella is limited by the amount of FlhF associated to the cell pole. Since *flhF* transcription is one of the targets for FleQ/FleN regulation, we propose that downregulation of FlhF synthesis is the primary mechanism by which FleN controls the number of flagella in these organisms. A second, posttranscriptional mechanism has been proposed for control of flagellar number. In *B. subtilis*, *C. jejuni*, and *Vibrio alginolyticus*, C-ring-associated FlhG may provoke FlhF dissociation by inducing its GTPase activity and the subsequent transition from an active, GTP bound dimer to an inactive, GDP-bound monomer (Bange *et al*., 2011; Altegoer *et al*., 2014; Schuhmacher *et al*., 2015; Gulbronson *et al*., 2016; Takekawa *et al*., 2016). Several lines of evidence suggest that this mechanism does not operate in *P. putida*. Firstly, FlhF stays bound to the cell poles for several cell cycles in FleN^+^ *P. putida* cells (**Fig. 4B and Supplemental Movie 2**). Secondly, the *P. putida* Δ*fimV* mutant, which fails to recruit FleN to the cell pole, is not hyperflagellated (**Fig. 8 A-C**), suggesting that polar location of FleN is not required for negative regulation of flagellar assembly. Finally, the activator helix, an N-terminal motif essential to stimulation of FlhF GTPase activity, is conspicuously absent in the *Pseudomonas* spp. FleN proteins (Schuhmacher *et al*., 2015).

Our observations suggest a second, previously undescribed, role for FleN and FimV in determining the timing of the onset of flagellar assembly. FleN and FimV prevent premature polar recruitment of FlhF and the early flagellar proteins FliF and FliM (**Fig. 7C-E**). Because FimV is required for polar FleN recruitment, we are inclined to believe that polar location of FleN is essential to this function, while FimV may or may not play an additional role other than facilitate the polar location of FleN. A previous report showed that deletion of *flhG* caused early polar recruitment of FlhF in *V. parahaemolyticus* (Arroyo-Pérez *et al*., 2021), but this phenomenon was not explored further. The biological relevance of the correct timing in the initiation of flagellar is as of now unclear and will be a subject of further investigation in the near future.

## ACKNOWLEDGMENTS

We wish to thank Carmen Beuzón, Javier Ruiz-Albert, Enrique Flores and Antonia Herrero for thoughtful advice and discussion, and Guadalupe Martín-Cabello and Isamar Moyano-Bravo for technical assistance.

## AUTHOR CONTRIBUTIONS

Marta Pulido-Sánchez: Data curation, formal analysis, investigation, methodology, writing - review and editing

Antonio Leal-Morales: Formal analysis, investigation, methodology

Aroa López-Sánchez: Conceptualization, methodology, project administration, writing - review and editing

Felipe Cava: Conceptualization, methodology, supervision

Fernando Govantes: Conceptualization, funding acquisition, methodology, project administration, resources, supervision, writing - original draft, writing - review and editing

