## Supplemental materials for "Spatial, temporal and numerical regulation of polar flagella assembly in *Pseudomonas putida*"

### 1. SUPPLEMENTAL MATERIALS AND METHODS

#### Plasmid and strain construction

**Construction of MRB97, MRB194, MRB175, MRB177 and MRB176.** To construct *P. putida* in-frame deletion mutants of the *fimV*, *fliF*, *fliM*, *fliC,* and *fliA* genes*,* upstream and downstream chromosomal regions flanking these genes were PCR-amplified using appropriate oligonucleotide pairs specified in **Supplemental Table S1**. Upstream and downstream *fimV* PCR products were assembled by overlap extension PCR and cloned *via* KpnI and BamHI restriction sites into the gene replacement vector pEMG that incorporates two I-SceI sites flanking the *lacZ* polylinker, yielding pMRB287. Upstream and downstream *fliF* and *fliC* PCR products were cleaved with XmaI and BamHI (upstream region) or BamHI and XbaI (downstream region), and three-way ligated into XmaI- and XbaI-digested pEMG, yielding pMRB356 and pMRB312, respectively. Upstream and downstream *fliM* PCR products were cleaved with BamHI and HindIII (upstream region) or HindIII and XbaI (downstream region), and three-way ligated into BamHI- and XbaI-digested pEMG, yielding pMRB314. Upstream and downstream *fliA* PCR products were ligated into SmaI-digested pEMG by Gibson assembly, yielding pMRB316. These plasmids were transferred by triparental mating to *P. putida* KT2442 harboring the pSW-I plasmid, expressing I-SceI from the 3-methyl benzoic acid-inducible *xylS*-P*m* system. Selection of integration and allelic replacement by homologous recombination-based repair of the chromosomal cleavage at the integrated I-SceI site was performed as described (Martínez-García & de Lorenzo, 2011) to generate Δ*fimV* (MRB97)*,* Δ*fliF* (MRB194)*,* Δ*fliM* (MRB175), Δ*fliC* (MRB177) and Δ*fliA* (MRB176) mutant strains*,* respectively. Loci deletions were verified by PCR. Curation of pSW-I plasmid was carried out by successive growth cycles in LB without antibiotics.

**Construction of pMRB437.** For FlgM-FLAG expression under the P*sal* promoter, a pMRB172-derivative was constructed by cloning the PCR-amplified *flgM* ORF carrying a C-terminal DYKDDDDK tag, into SpeI-PstI digested pMRB172, yielding pMRB437.

**Construction of pMRB189, pMRB236, pMRB238 and pMRB375.** For the construction of pMRB189 and pMRB238 mini-Tn*7* delivery vectors for GFP translational fusions to proteins at their C-terminus and N-terminus, expressed from the salicylate-inducible P*sal* promoter, the *gfp*mut3 gene from pMRB1 was PCR-amplified with oligonucleotide pairs specified in **Supplemental Table S1** and cloned into SpeI- and PstI-digested pMRB172. For FimV-GFP expression from the P*sal* promoter, a pMRB189 derivative was constructed by cloning the PCR-amplified *fimV* ORFs lacking the stop codon into SpeI- and BamHI-digested pMRB189, yielding pMRB236. For GFP-FleN expression from the P*sal* promoter, a pMRB238 derivative was constructed by cloning the PCR-amplified *fleN* ORF into SacI- and BamHI-digested pMRB238, yielding pMRB375.

**Construction of pMRB187, pMRB200, pMRB310 pMRB374, and pMRB451.** For the construction of pMRB187, a mini-Tn*7* delivery vector for C-terminal GFP translational fusions, the *gfp*-mut3 gene from pMRB1 was PCR-amplified and cloned into SpeI- and PstI-digested pUC18Sfi-miniTn*7*BB-Gm. For FlhF-GFP expression under its own P*flhF* promoter, a P*flhF-flhF* chromosomal region was PCR-amplified and cloned into SpeI- and BamHI-digested pMRB187, yielding pMRB200. For FliF-GFP, FliM-GFP and FimV-GFP expression under their operon promoters P*fliE,* P*fliL* and P*asd,* respectively, a Gibson assembly strategy was followed for cloning into pMRB187, adding some in-frame start and end nucleotides of *fliE*, *fliL* and *asd* downstream the P*fliE*, P*fliL* and P*asd* promoters to provide transcriptional and translational context for *fliF*, *fliM* and *fimV* expression. A fragment containing the P*fliE*, P*fliL* or *Pasd* promoter regions and the first 54 bp of *fliE*, 69 bp of *fliL* or 72 bp of *asd*, and a second fragment containing the last 57 bp of *fliE* or *fliL*, or 81 bp of *asd* followed by the *fliF*, *fliM* or *fimV* coding regions were PCR-amplified. PCR products corresponding to P*fliE-fliF* and *Pasd-fimV* were ligated with SmaI-digested pMRB187 by Gibson assembly, yielding pMRB374 and pMRB451. PCR products corresponding to P*fliL-fliM* were ligated by Gibson assembly with PCR-amplified pMRB187, yielding pMRB310.

**Construction of pMRB302.** For the expression of FliC$^{S267C}$ under its own P*fliC* promoter, site-directed mutagenesis PCR was performed for serine-to-cysteine replacement with appropriate oligonucleotides (Hintsche *et al*., 2022), and overlap extension PCR product holding P*fliC-fliC*$^{S267C}$ was cloned into XbaI- and SpeI-digested pUC18Sfi-miniTn*7*BB-Gm, yielding pMRB302.

### 2. SUPPLEMENTAL TABLE

**Supplemental Table S1. Bacterial strains, plasmids and oligonucleotides used in this work.** Cm: chloramphenicol. Rif: rifampicin. Km: kanamycin. Ap: ampicillin. Gm: gentamycin.

| Bacterial strain | Genotype/phenotype | Reference/source |
|---|---|---|
| ***E. coli*** | | |
| DH5α | Φ80d*lacZ*ΔM15 Δ(*lacZYA-argF*)U169 *recA*1 *endA*1 *hsdR*17 ($r_k^-$ $m_k^+$) *supE*44 *thi*-1 *gyrA relA*1 | Hanahan, 1983 |
| DH5α λpir | DH5α with lysogenic phage λ-pir, host for R6K replication origin plasmids | Víctor de Lorenzo |
| ***P. putida*** | | |
| KT2442 | mt-2 *hsdR*1 ($r^-$ $m^+$). Cm$^r$ Rif$^r$. | Franklin *et al*., 1981 |
| MRB175 | KT2442 Δ*fliM*. Cm$^r$ Rif$^r$. | This work |
| MRB176 | KT2442 Δ*fliA*. Cm$^r$ Rif$^r$. | This work |
| MRB177 | KT2442 Δ*fliC.* Cm$^r$ Rif$^r$. | This work |
| MRB194 | KT2442 Δ*fliF*. Cm$^r$ Rif$^r$. | This work |
| MRB52 | KT2442 Δ*fleQ*. Cm$^r$ Rif$^r$. | Navarrete *et al*., 2019 |
| MRB62 | KT2442 *fliP*::miniTn5-Km. Cm$^r$ Rif$^r$ Km$^r$ | López-Sánchez *et al*., 2016 |
| MRB69 | KT2442 Δ*flhF*. Cm$^r$ Rif$^r$. | Navarrete *et al*., 2019 |
| MRB71 | KT2442 Δ*fleN*. Cm$^r$ Rif$^r$. | Navarrete *et al*., 2019 |
| MRB97 | KT2442 Δ*fimV*. Cm$^r$ Rif$^r$. | This work |

| Plasmid | Genotype/phenotype | Reference/source |
|---|---|---|
| pEMG | pJP5603 bearing a *lacZ*α polylinker with two flanking I-SceI sites. R6K, Mob$^+$, Km$^r$ | Martínez-García & de Lorenzo, 2011 |
| pMRB1 | pBBR1-MCS4-derived broad host-range *gfp*mut3::*lac*Z transcriptional fusion vector. pBBR1, Mob$^+$, Ap$^r$ | Jiménez-Fernández *et al*., 2015 |
| pMRB172 | pUC18Sfi-miniTn*7*BB-Gm-based delivery plasmid for miniTn*7*BB-Gm [*nahR*-P*sal*]. Ap$^r$ Gm$^r$ | Leal-Morales *et al*., 2022 |
| pMRB187 | pUC18Sfi-miniTn*7*BB-Gm-based delivery plasmid for C-terminal *gfp*-mut3 translational fusions. Ap$^r$ Gm$^r$ | This work |
| pMRB189 | pMRB172-derived delivery plasmid for C-terminal *gfp*-mut3 translational fusions. Ap$^r$ Gm$^r$ | This work |
| pMRB200 | pMRB187-derived vector containing P*flhF-flhF* translationally fused to *gfp*mut3 at its C-terminus. Ap$^r$ Gm$^r$ | This work |
| pMRB236 | pMRB189-derived vector containing a RBS site and *fimV* translationally fused to *gfp*mut3 at its C-terminus. Ap$^r$ Gm$^r$ | This work |
| pMRB238 | pMRB172-derived vector containing a RBS site and *gfp*mut3 N-terminal translational fusions Ap$^r$ Gm$^r$. | This work |
| pMRB250 | pMRB3-derived vector containing a *gfp*mut3::*lacZ* transcriptional fusion to the P*flgF* promoter. Ap$^r$ | Leal-Morales *et al*., 2022 |
| pMRB259 | pMRB3-derived vector containing a *gfp*mut3::*lacZ* transcriptional fusion to the P*fliE* promoter. Ap$^r$ | Leal-Morales *et al*., 2022 |
| pMRB260 | pMRB3-derived vector containing a *gfp*mut3::*lacZ* transcriptional fusion to the P*fliL* promoter. Ap$^r$ | Leal-Morales *et al*., 2022 |
| pMRB265 | pMRB3-derived vector containing a *gfp*mut3::*lacZ* transcriptional fusion to the P*flhF* promoter. Ap$^r$ | Leal-Morales *et al*., 2022 |
| pMRB270 | pMRB3-derived vector containing a *gfp*mut3::*lacZ* transcriptional fusion to the P*fliC* promoter. Ap$^r$ | Jiménez-Fernández *et a*l., 2016 |
| pMRB271 | pMRB3-derived vector containing a *gfp*mut3::*lacZ* transcriptional fusion to the P*motA* promoter. Ap$^r$ | Jiménez-Fernández *et al*., 2016 |
| pMRB272 | pMRB3-derived vector containing a *gfp*mut3::*lacZ* transcriptional fusion to the P*flgB* promoter. Ap$^r$ | Jiménez-Fernández *et al*., 2016 |
| pMRB276 | pMRB3-derived vector containing a *gfp*mut3::*lacZ* transcriptional fusion to the P*cheA* promoter. Ap$^r$ | Jiménez-Fernández *et al*., 2016 |
| pMRB287 | pEMG bearing a 1600 bp insert with upstream and downstream flanking regions of *fimV*. Km$^r$ | This work |
| pMRB3 | pMRB1-derived vector containing the Gateway conversion cassette *attR2-ccdB*-Cm$^r$-*attR1*. Ap$^r$ Cm$^r$ | Jiménez-Fernández *et al*., 2015 |
| pMRB302 | pUC18Sfi-mini*Tn7*BB-Gm-derived vector containing P*fliC-fliC*$^{S267C}$. Ap$^r$ Gm$^r$ | This work |
| pMRB310 | pMRB187-derived vector containing P*fliL-fliM* translationally fused to *gfp*mut3 at its C-terminus. Ap$^r$ Gm$^r$ | This work |
| pMRB312 | pEMG bearing a 897 bp insert with upstream and downstream flanking regions of *fliC*. Km$^r$ | This work |
| pMRB314 | pEMG bearing a 1005 bp insert with upstream and downstream flanking regions of *fliM*. Km$^r$ | This work |

| | | |
|---|---|---|
| pMRB316 | pEMG bearing a 1002 bp insert with upstream and downstream flanking regions of *fliA*. Km$^r$ | This work |
| pMRB356 | pEMG bearing a 1012 bp insert with upstream and downstream flanking regions of *fliF*. Km$^r$ | This work |
| pMRB374 | pMRB187-derived vector containing P*fliE-fliF* translationally fused to *gfp*mut3 at its C-terminus. Ap$^r$ Gm$^r$ | This work |
| pMRB375 | pMRB238-derived vector containing *fleN* translationally fused to *gfp*mut3 at its N-terminus. Ap$^r$ Gm$^r$ | This work |
| pMRB437 | pMRB172-derived vector containing a RBS site and a C-terminal *flgM*-FLAG tag fusion. Ap$^r$ Gm$^r$ | This work |
| pMRB451 | pMRB187-derived vector containing P*asd-fimV* translationally fused to *gfp*mut3 at its C-terminus. Ap$^r$ Gm$^r$ | This work |
| pRK2013 | Helper plasmid for triparental mating. ColE1, Mob$^+$, Km$^r$ | Figurski & Helinski, 1979 |
| pSW-I | Plasmid expressing I-SceI from *xylS*-P*m*. RK2, Mob$^+$, Ap$^r$ | Wong & Mekalanos, 2000 |
| pTNS2 | Helper plasmid expressing the *Tn*7 transposase. R6K, Mob$^+$, Ap$^r$ | Choi *et al*., 2005 |
| pUC18Sfi-miniTn*7*BB-Gm | pUC18SfiI-based delivery plasmid for the synthetic minitansposon mini*Tn*7BB-Gm. Ap$^r$ Gm$^r$ | Jiménez-Fernández *et al*., 2015 |

| **Oligonucleotide** | **Sequence (5' to 3')** | **Use** |
|---|---|---|
| Asd5`-asd3`_fwd | GCCTTGGTGCAGATCCTCCTAACCACCGACAACGTGC | Amplification of last 81 bp of *asd* and *fimV* ORF for Gibson assembly |
| Asd5'-asd3'_rev | GCACGTTGTCGGTGGTTAGGAGGATCTGCACCAAGGC | Amplification of P*asd* and first 72 bp of *asd* for Gibson assembly |
| FimV-BamHI_rev | ACGTGGATCCGACCAGCCGGGAGAGCAT | *fimV* ORF amplification |
| FimV-DOWN-BamHI_rev | ACGTGGATCCCAGCACACCGGCAATGTTAC | *fimV* downstream region amplification |
| FimV-DOWN-DOWN_rev | AGGCAACCGGACGCTC | *fimV* chromosomal deletion verification |
| FimV-qPCR_fwd | CACAGAATCAGCCGCTGGAC | qRT-PCR of *fimV* ORF |
| FimV-qPCR_rev | CCTTGCTGAACTCTTCCGGTG | qRT-PCR of *fimV* ORF |
| FimV_rev | GACCAGCCGGGAGAGCAT | Amplification of last 81 bp of *asd* and *fimV* ORF for Gibson assembly |
| FimV-SpeI-SD_fwd | ACGTACTAGTGAAAGAGGAGAAATACTAGATGCTTCGAATTCGCAAACTG | *fimV* ORF amplification |
| FimV-UP-DOWN-overlap_fwd | GATTCATACAAGGGAAGAGGTCTGCAAGCAGGTCAGGC | *fimV* downstream region amplification |
| FimV-UP-DOWN-overlap_rev | GCCTGACCTGCTTGCAGACCTCTTCCCTTGTATGAATC | *fimV* upstream region amplification |
| FimV-UP-SacI-KpnI_fwd | ACGTGAGCTCGGTACCGTCATCGACCTGTCCGGC | *fimV* upstream region amplification |
| FimV-UP-UP_fwd | CGTCAGCCGCAGCTTTG | *fimV* chromosomal deletion verification |
| FleN-BamHI_rev | AGCTGGATCCTCATAGTACGGGTCCCGC | *fleN* ORF amplification |
| FleN-SacI_fwd | AGCTGAGCTCGGTAGCATGCATCCCGTA | *fleN* ORF amplification |
| FlgM-FLAG-PstI_rev | CATGCTGCAGTTACTTGTCGTCATCGTCTTTGTAGTCGCGCTGGGCTTCGAAATC | *flgM* ORF amplification with FLAG tag |
| FlgM-XbaI_fwd | CAGTTCTAGAGAAAGAGGAGAAAATGGTCATCGACTTCAGTCGTTTG | *flgM* ORF amplification with FLAG tag |
| FlhF-BamHI_rev | ACGTGGATCCACCCGCTCGCCGTGG | *flhF* ORF amplification |
| FliA-DOWN-DOWN_rev | ATGTTCCAGTCGGTCACC | *fliA* chromosomal deletion verification |
| FliA-DOWN-GA_fwd | CCCACCAGCGCGGGACCCGTACTATGCATGAGGATGGCGCCAGTGGC | *fliA* downstream region amplification |
| FliA-DOWN-GA_rev | CTGCAGGTCGACTCTAGAGGATCCCCGTAGTGACCGTTTTCCAGCAT | *fliA* downstream region amplification |
| FliA-UP-GA_fwd | GGCCAGTGAATTCGAGCTCGGTACCCTTCAGCGAGATCGGCGACAAC | *fliA* upstream region amplification |
| FliA-UP-GA_rev | ACCTGGCCACTGGCGCCATCCTCATGCATAGTACGGGTCCCGCGCTG | *fliA* upstream region amplification |
| FliA-UP-UP_fwd | AGAGCATGGTGCATCTGG | *fliA* chromosomal deletion verification |
| FliC-DOWN-BamHI_fwd | GATCGGATCCTAATTCGATTCTGTGATGAATCG | *fliC* downstream region amplification |
| FliC-DOWN-DOWN_rev | GACACAATCCGAGTCAAGAC | *fliC* chromosomal deletion verification |
| FliC-DOWN-XbaI_rev | GATCTCTAGACATCACCCTCCGCCAGA | *fliC* downstream region amplification |
| FliC-mutSer267_fwd | TGTTACCGCTTGCATTGACAG | *fliC* ORF amplification with S267C replacement |
| FliC-mutSer267_rev | CTGTCAATGCAAGCGGTAACA | *fliC* ORF amplification with S267C replacement |
| FliC-SpeI_rev | ACGTACTAGTTTAGCCGAGCAGCTTCAG | *fliC* ORF amplification |
| FliC-UP-BamHI_rev | GATCGGATCCCATGACGAATTCCTCGTTGTA | *fliC* upstream region amplification |

| | | |
|---|---|---|
| FliC-UP-UP_fwd | AAGGACATGCTGGAAACC | *fliC* chromosomal deletion verification |
| FliC-UP-XmaI_fwd | GATCCCCGGGCTGCTGCTGCAGAAGCAC | *fliC* upstream region amplification |
| FliE5'-fliE3'-GA_fwd | TGATGTTGGACATGCGGGCCATGCAGCAGGTACGCAACAAGCTGGTC | Amplification of last 57 bp of *fliE* and *fliF* ORF for Gibson assembly |
| FliE5'-fliE3'-GA_rev | GCCTGGACCAGCTTGTTGCGTACCTGCTGCATGGCCCGCATGTC | Amplification of P*fliE* and first 54 bp of *fliE* for Gibson assembly |
| FliF-DOWN-BamHI_fwd | GATCGGATCCGCCGATGAGTGATAACCG | *fliF* downstream region amplification |
| FliF-DOWN-DOWN_rev | CTGGTTCAGCTCCTTCAGC | *fliF* chromosomal deletion verification |
| FliF-DOWN-XbaI_rev | GATCTCTAGAGTGTTCAGCGATGACACG | *fliF* downstream region amplification |
| FliF-pMRB187-GA_rev | AGTTCTTCTCCTTTACTGGATCCCCCCTCATCGGCGTTGATCCA | Amplification of last 57 bp of *fliE* and *fliF* ORF for Gibson assembly |
| FliF-UP-BamHI_rev | GATCGGATCCCATGGGTTACATCCGTCCTC | *fliF* upstream region amplification |
| FliF-UP-UP_fwd | CCAGGTCTTCAGCTCCG | *fliF* chromosomal deletion verification |
| FliF-UP-XmaI_fwd | GATCCCCGGGCCTATCGCGACGCAAGG | *fliF* upstream region amplification |
| FliL5'-fliL3'-GA_fwd | AAACTCAAGCTGATCCTGCTGGCTGTCGAAGTCGGCAAGCCGGTCATC | Amplification of last 57 bp of *fliL* and *fliM* ORF for Gibson assembly |
| FliL5'-fliL3'-GA_rev | CTGGTCGATGACCGGCTTGCCGACTTCGACAGCCAGCAGGATCAGCTT | Amplification of P*fliL* and first 69 bp of *fliL* for Gibson assembly |
| FliM-DOWN-DOWN_rev | CACTCAGCGCAGCTTCTT | *fliM* chromosomal deletion verification |
| FliM-DOWN-HindIII_fwd | GATCAAGCTTGACCCGATCGAACGCC | *fliM* downstream region amplification |
| FliM-DOWN-XbaI_rev | GATCTCTAGAGGCTGATCACGTCGGTCA | *fliM* downstream region amplification |
| FliM-pMRB187-GA_fwd | CAGATCATCGACCCGATCGAACGCCGCACTAGTGAGCTCCCCGGGGGA | pMRB187 amplification for Gibson assembly |
| FliM-pMRB187-GA_rev | ACTGGATCCCCCGGGGAGCTCACTAGTGCGGCGTTCGATCGGGTCGAT | Amplification of last 57 bp of *fliL* and *fliM* ORF for Gibson assembly |
| FliM-UP-BamHI_fwd | GATCGGATCCGAAGCAGTCAAAGACCCC | *fliM* upstream region amplification |
| FliM-UP-HindIII_rev | GATCAAGCTTCATTGCGGACTCCTACTG | *fliM* upstream region amplification |
| FliM-UP-UP_fwd | TGGACGACGGATTATTGG | *fliM* chromosomal deletion verification |
| GFP-SacI-SmaI-BamHI-PstI_rev | AGCTCTGCAGGGATCCCCCGGGGAGCTCTTTGTATAGTTCATCCATGCCATGT | Amplification of *gfp*mut3 for pMRB238 construction |
| Pasd_fwd | CAAGGTGCTCAAAGCTAAGGCC | Amplification of P*asd* and first 72 bp of *asd* for Gibson assembly |
| pMRB187-pfliE-GA_fwd | GCTTCTAGAGTACTAGTGAGCTCCCCGCGTCGTCACGAGTACCAGATGAT | Amplification of P*fliE* and first 54 bp of *fliE* for Gibson assembly |
| pMRB187-pfliL-GA_fwd | TCTGGAATTCGCGGCCGCTTCTAGAGTTCAAGTCGGCTGATATCCAAC | Amplification of P*fliL* and first 69 bp of *fliL* for Gibson assembly |
| pMRB187-pfliL-GA_rev | GCTCCAGTTGGATATCAGCCGACTTGAACTCTAGAAGCGGCCGCGAAT | pMRB187 amplification for Gibson assembly |
| PflhF-flhF-SpeI_fwd | AGCTACTAGTGCTCTGAAGACATGGGCAAGCT | P*flhF* promoter and *flhF* ORF amplification |
| PfliC-fliC-XbaI_fwd | ACGTTCTAGAAAGCACATGCTCGATGGC | P*fliC* promoter and *fliC* ORF amplification |
| PstI-GFP_rev | ACGTCTGCAGTTATTATTATTTGTATAGTTCATCCATGCCA | Amplification of *gfp*mut3 for pMRB187 and pMRB189 construction |
| SpeI-GFP_fwd | AGCTACTAGTGAAAGAGGAGAAATACTAGATGAGTAAAGGAGAAGAACTTTTCACTG | Amplification of *gfp*mut3 for pMRB238 construction |
| SpeI-SacI-SmaI-BamHI-GFP_fwd | ACGTACTAGTGAGCTCCCCGGGGGATCCAGTAAAGGAGAAGAACTTTTCACTGG | Amplification of *gfp*mut3 for pMRB187 and pMRB189 construction |
| Tn7-GlmS | AATCTGGCCAAGTCGGTGAC | Confirmation of miniTn7-derivatives integration |
| Tn7-R109 | CAGCATAACTGGACTGATTTCAG | Confirmation of miniTn7-derivatives integration |

### 3. SUPPLEMENTAL FIGURES

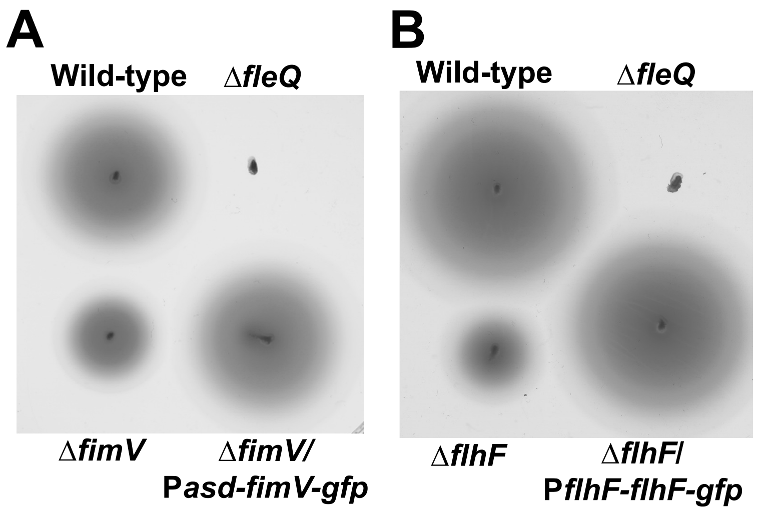


**Figure S1. Complementation of the Δ*fimV* and Δ*flhF* mutants.** Soft agar-based swimming assays showing the complementation of the Δ*fimV* mutant with P*asd-fimV-gfp* **(A)** or the Δ*flhF* mutant with P*flhF-flhF-gfp* **(B)**. The wild-type strain and the Δ*fleQ* mutant assayed in the same plate were used as positive and negative controls. The picture shows a representative swim plate out of at least three separate replicates.

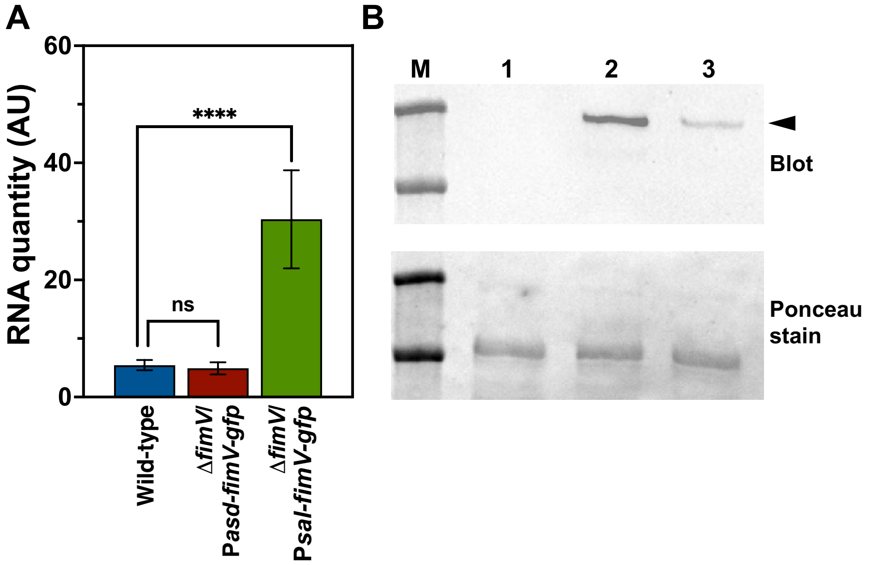


**Figure S2. Expression of FimV. A.** qRT-PCR quantification of *fimV* expression in exponential phase cultures of the wild-type strain, and the Δ*fimV* mutant bearing the P*asd-fimV-gfp* or P*sal-fimV-gfp* tranposon. Columns and error bars represent averages and standard deviations of at least three separate replicates. Stars denote *p*-values of the two-tailed Student's T test not assuming equal variance (*=$p$<0.05; **= $p$<0.01; ***= $p$<0.001; ****= $p$<0.0001). **B.** Western blot analysis (top) of FimV-GFP in the wild-type strain (lane 1) or the wild-type bearing the P*sal-fimV-gfp* (lane 2) or P*asd-fimV-gfp* (lane 3) transposon using anti-GFP antiserum- The arrowhead indicates the FimV-GFP fusion protein. The size marker lane (M) shows 250 and 150 kDa bands. The Ponceau stained-membrane is shown (bottom) as protein loading control.

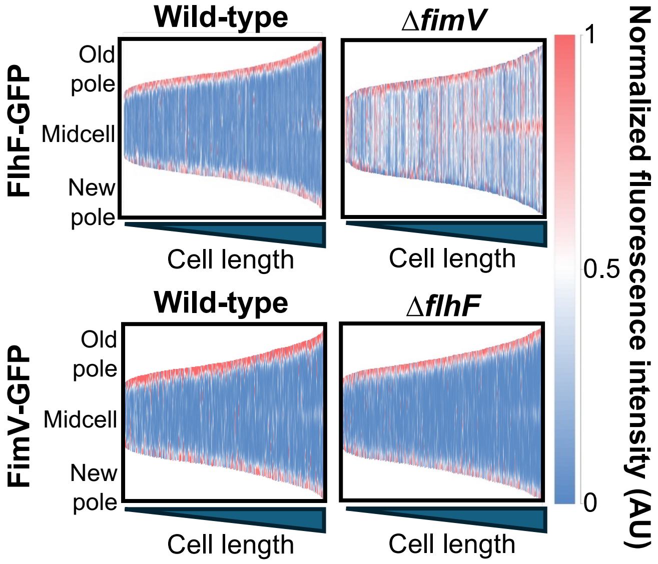


**Figure S3. FlhF-GFP and FimV-GFP fluorescence demographic maps of length-sorted cells.** Vertical slabs represent normalized FlhF-GFP (top) of FimV-GFP (bottom) fluorescence intensity along the medial cell axis of 500 length-sorted wild-type (left), Δ*fimV* (top right) or Δ*flhF* (bottom right) cells. Intensity is displayed in a blue-white-red scale with blue representing low, red representing high and white representing intermediate values.

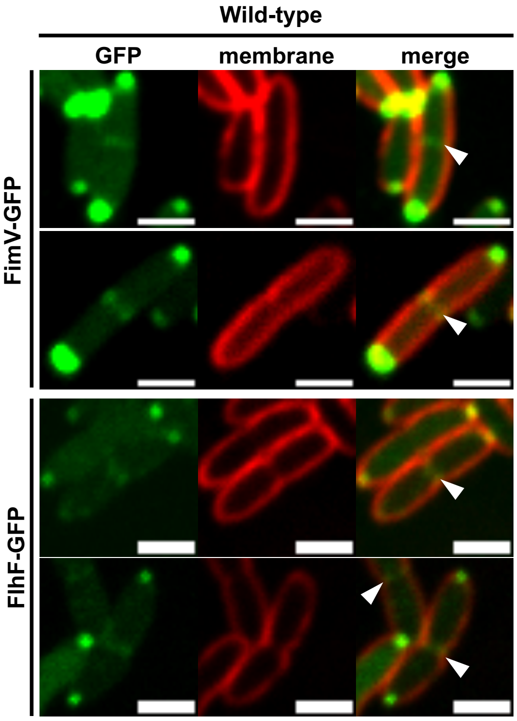


**Figure S4. FimV-GFP and FliF-GFP midcell bands.** Confocal microscopy images of wild-type cells expressing P*asd-fimV-gfp* or P*flhF-flhF-gfp* (green). FM$^{TM}$ 4-64 was used as membrane stain (red). Images show the maxima projections of seven Z-sections of the green channel (GFP), the red channel showing the cell contour at the focal plane (membrane) and the composite of both channels (merge). Arrowheads indicate fluorescent midcell bands. Scale bar: 2 µm.

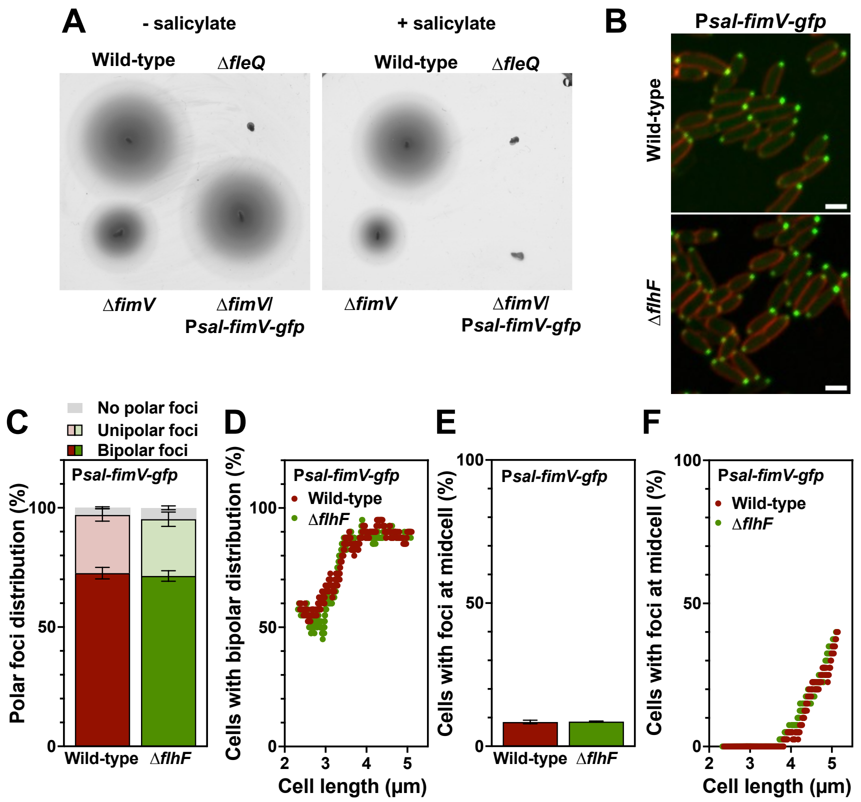


**Figure S5. Complementation and intracellular location of FimV expressed under P*sal*. A.** Soft agar-based swimming assays showing the complementation of the Δ*fimV* mutant with P*sal-fimV-gfp* in the absence (left) or the presence (right) of 2 mM salicylate. The wild-type strain and the Δ*fleQ* mutant assayed in the same plate were used as positive and negative controls. The picture shows a representative swim plate out of at least three separate replicates. **B.** Confocal microscopy images of wild-type and Δ*flhF* cells expressing P*sal-fimV-gfp* in the absence of salicylate. FM$^{TM}$ 4-64 was used as membrane stain (red). Images are shown as the maxima projections of seven Z-sections of the green channel, merged with the red channel showing the cell contour at the focal plane. Scale bar: 2 μm. **C.** Frequency of wild-type and Δ*flhF* cells bearing bipolar, unipolar or no polar FimV-GFP foci. **D.** Frequency of wild-type and Δ*flhF* cells displaying bipolar distribution of FimV-GFP foci *vs.* cell length (n=500 cells). **E and F.** Frequency of wild-type and Δ*flhF* cells displaying midcell FimV-GFP foci **(E)** and the same parameter plotted against cell length **(F)** (n=500 cells). Columns and error bars represent averages and standard deviations of at least three separate replicates. Stars denote *p*-values of the two-tailed Student's T test not assuming equal variance (*=$p$<0.05; **= $p$<0.01; ***= $p$<0.001; ****= $p$<0.0001).

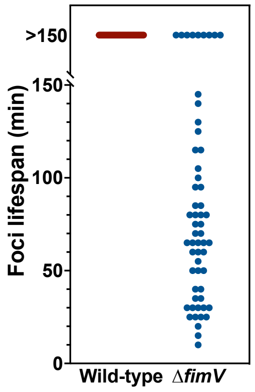


**Figure S6. Lifespan of polar FlhF-GFP foci.** Points represent the lifespan measurements of individual foci calculated from images acquired in five minute intervals during the course of five hours. Lifespan values were registered for a maximum of 150 minutes.

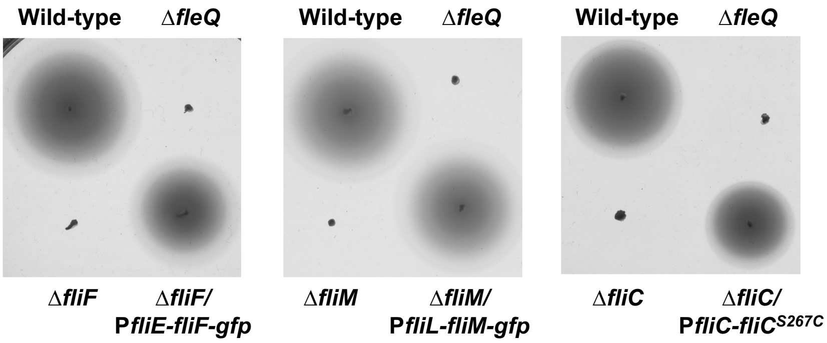


**Figure S7. Complementation of the Δ*fliF,* Δ*fliM* and Δ*fliC* mutants.** Soft agar-based swimming assays showing the complementation of the Δ*fliF,* Δ*fliM* and Δ*fliC* mutants with P*fliE-fliF-gfp,* P*fliL-fliM-gfp* or P*fliC-fliC*$^{S267C}$, respectively. The wild-type strain and the Δ*fleQ* mutant assayed in the same plate were used as positive and negative controls. The picture shows a representative swim plate out of at least three separate replicates.

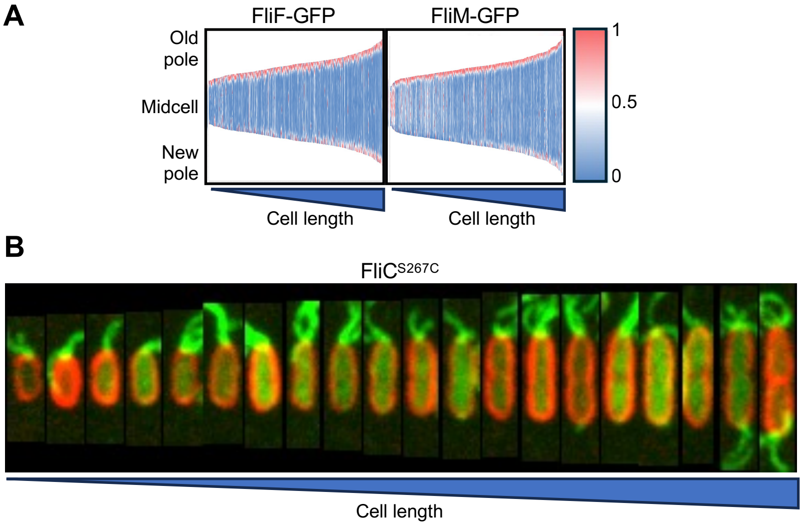


**Figure S8. Recruitment dynamics of the flagellar structural proteins to the new cell pole. A.** Fluorescence demographic maps of wild-type length-sorted cells expressing P*fliE-fliF-gfp* or P*fliL-fliM-gfp* (n=500 cells). Vertical slabs represent normalized fluorescence intensity along the medial cell axis. Intensity is displayed in a blue-white-red scale with blue representing low, red representing high and white representing intermediate values. **B.** Virtual kymograph of wild-type length-sorted cells expressing P*fliC-fliC*$^{S267C}$ (green, n=20 cells). Alexa Fluor$^{TM}$ 488 $C_5$ maleimide was used for FliC$^{S267C}$ filament staining. FM$^{TM}$ 4-64 was used as membrane stain (red). Images are shown as the maxima projections of seven Z-sections of the green channel, merged with the red channel showing the cell contour at the focal plane.

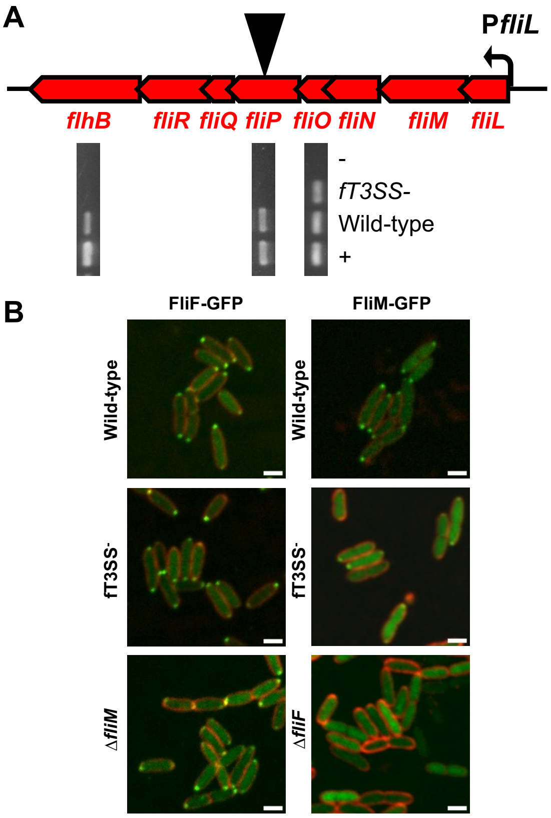


**Figure S9. Intracellular location of early basal body components. A.** RT-PCR analysis of the fT3SS⁻ mutant. Cartoon of the *fliLMNOPQR-flhB* operon showing the location of the miniTn*5*-Km transposon insertion as a black inverted triangle (top), and agarose gel images showing RT-PCR amplification products using cDNA from the wild-type strain or the fT3SS⁻ mutant, chromosomal DNA of the wild-type strain (+), or no template (-), and primers specific for *flhB*, *fliP* or *fliO* (bottom). **B.** Confocal microscopy images of wild-type, fT3SS⁻ and Δ*fliM* (left) or Δ*fliF* (right) cells expressing P*fliE-fliF-gfp* (left) or P*fliL-fliM-gfp* (right). FM™ 4-64 was used as membrane stain (red). Images are shown as the maxima projections of seven *Z*-sections of the green channel, merged with the red channel showing the cell contour at the focal plane. Scale bar: 2 µm.

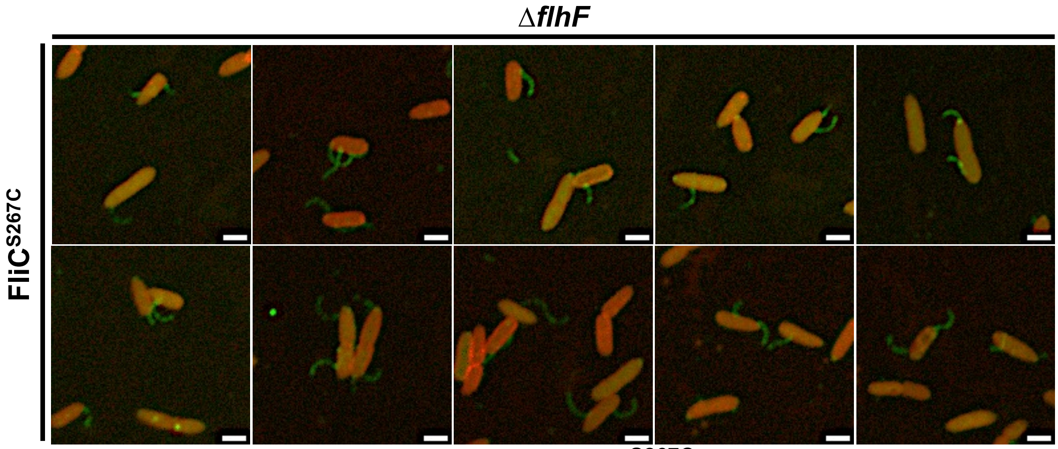


**Figure S10. Confocal microscopy images of FliC$^{S267C}$ filaments in the Δ*flhF* mutant.** Ten representative images of Δ*flhF* cells are shown. Alexa Fluor™ 488 $C_5$ maleimide was used for FliC$^{S267C}$ filament staining. FM™ 4-64 was used as membrane stain (red). Images are shown as the maxima projections of seven Z-sections of the green channel, merged with the red channel showing the cell contour at the focal plane. Scale bar: 2 µm.

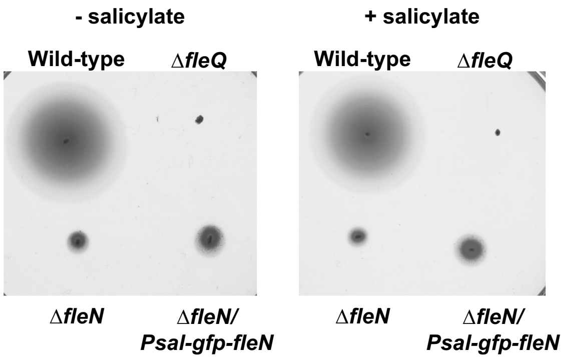


**Figure S11. Complementation of the Δ*fleN* mutant.** Soft agar-based swimming assays showing the complementation of the Δ*fleN* mutant with P*sal-gfp-fleN* in the absence (left) or the presence (right) of 2 mM salicylate. The wild-type strain and the Δ*fleQ* mutant assayed in the same plate were used as positive and negative controls. The picture shows a representative swim plate out of at least three separate replicates.

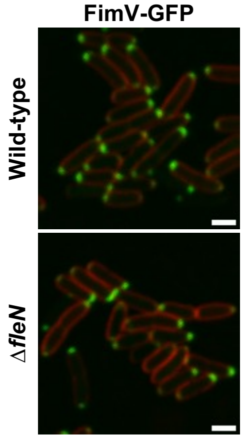


**Figure S12. Intracellular location of FimV in the Δ*fleN* mutant.** Confocal microscopy images of wild-type and Δ*fleN* cells expressing P*asd-fimV-gfp*. FM$^{TM}$ 4-64 was used as membrane stain (red). Images are shown as the maxima projections of seven Z-sections of the green channel, merged with the red channel showing the cell contour at the focal plane. Scale bar: 2 µm.

### 4. SUPPLEMENTAL MOVIES

**Supplemental movie S1.** Fluorescence microscopy time-lapse of wild-type cells expressing P*sal-fimV-gfp*. Images were captured in 5-minute intervals for 300 minutes

**Supplemental movie S2.** Fluorescence microscopy time-lapse of wild-type cells expressing P*flhF-flhF-gfp*. Images were captured in 5-minute intervals for 300 minutes

**Supplemental movie S3.** Fluorescence microscopy time-lapse of Δ*fimV* cells expressing P*flhF-flhF-gfp*. Images were captured in 5-minute intervals for 300 minutes
